# Development of a qPCR assay and tremabiome deep amplicon sequencing method for differentiation of fluke species in livestock

**DOI:** 10.1101/2025.07.31.667929

**Authors:** Muhammad Abbas, Kezia Kozel, Olukayode Daramola, Nick Selemetas, Qasim Ali, Shoaib Ashraf, Isah Ibrahim, Inaki Deza-Cruz, Sai Fingerhood, Mark W. Robinson, Eric R Morgan, Umer Chaudhry, Martha Betson

## Abstract

**Background:** Trematode parasites, or flukes, are a significant economic threat to ruminant production worldwide. Traditional diagnostic methods rely on egg sedimentation from faeces, a time-consuming methodology lacking sensitivity and specificity. This study aimed to develop and validate two diagnostic methods: firstly, qPCR for accurate identification of *Fasciola* spp, and secondly, tremabiome, deep amplicon sequencing technique for identifying fluke species using faecal egg DNA.

**Methodology:** To detect fluke infection primers targeting mitochondrial DNA were repurposed to develop a SYBR Green qPCR diagnostic. For the identification of fluke species, a tremabiome approach was developed. A reference sequence library and taxonomy file were generated for 21 fluke species, enabling species sequence read separation and extracting amplicon sequence variants (ASVs). To validate the qPCR and tremabiome approach, 402 faecal samples were collected from cattle and sheep across the UK. Fluke eggs were isolated by sedimentation, detected by microscopy and qPCR, and tremabiome used to identify fluke eggs to species level.

**Results:** qPCR demonstrated high analytical sensitivity, detecting *Fasciola hepatica* DNA down to 19.2fg and *F. gigantica* down to 6.4fg, with no cross-amplification of other flukes. Tremabiome was able to detect as few as five *F. hepatica* and *Calicophoron daubneyi* eggs and identify mixed infections. High levels of co-infection (14.4%) of *F. hepatica* and *C. daubneyi* were observed in faecal samples, followed by single infections with *C. daubneyi* (12.6%) and *F. hepatica* (3.2%). Notably, tremabiome detected *F. hepatica* in 20 samples missed by qPCR. Data analysis identified 55 and 32 ASVs for *F. hepatica* and *C. daubneyi*, respectively, with phylogenetic clustering within their respective clades.

**Conclusion:** This study developed qPCR assay for *Fasciola* detection and validated a tremabiome deep amplicon sequencing for fluke species differentiation. These approaches have improved capacity to identify fluke species compared to microscopy and are valuable tools for enhancing fasciolosis surveillance and control.

**Author Summary:** Flukes are flatworm parasites that cause disease domestic and wild animals and humans. The main species infecting cattle and sheep globally are the liver flukes *F. hepatica* and *F. gigantica*, with other species including the rumen fluke *Calicophoron daubneyi* locally important or emerging. Infections result in serious economic losses. The traditional method of diagnosing fluke infection involves observation of eggs in faecal samples under the microscope, but this can be time-consuming and error prone, since the eggs of different species often look similar. In this study, we developed and validated two methods to improve detection: qPCR, a sensitive DNA-based test to identify *Fasciola* infections, and tremabiome, a DNA sequencing technique that can accurately differentiate between different fluke species. We tested these methods using faecal samples collected from cattle and sheep across the UK. The qPCR could detect small amounts of *Fasciola* DNA, while tremabiome was more sensitive, identifying different fluke species from as few as five eggs. Our study found that co-infections of *F. hepatica* and *C. daubneyi* are common in the UK. The approaches we have developed could be valuable tools for to improve fluke diagnosis and enable better control of this important parasitic disease.

## Introduction

Trematode parasites, or flukes, are widespread globally and include several species that cause serious disease in animals and humans. Fasciolosis is a neglected foodborne tropical disease caused by the zoonotic flukes *Fasciola hepatica* and *Fasciola gigantica*. Unlike other neglected tropical diseases, *Fasciola* infections in humans and animals have a broad reach globally, being found in more than 75 countries, with 2.4 million people infected, and millions more at risk [1]. The prevalence in livestock is less well known, however, a recent meta-analysis suggests the global prevalence of fasciolosis in cattle and sheep across continents ranges from 12-97% and 9-58% respectively across continents [2].

Ensuring food security is increasingly challenging with a growing global population. In 2020, the agri-food sector contributed 115 billion GBP, making up 6.0% of the UK economy [3], with other national economies considerably more dependent on farming. Recent global estimates indicate that fasciolosis may cost annual losses in animal productivity exceeding US$3.2 billion [4,5]. In the UK, fasciolosis prevails in ruminants, costing the cattle industry 13-40 million GBP yearly, reducing dairy farms’ net profit by 12% and beef farms’ by 6% [6]. *Fasciola* infections can lead to delayed animal slaughter [7], and condemnation of damaged livers [8]. According to the Food Standards Agency in 2014, 22% of British cattle livers are condemned due to fluke [9]. Losses to fasciolosis are widespread; for example, Australia faces one of the highest disease burdens, with estimated annual losses reaching approximately 129 million (range 38–193 million) AUD annually [10]. Infected animals suffer reduced weight, anaemia, reduced milk yield and fat content [8], lower reproduction, and higher mortality [11].

Flukes other than *Fasciola* spp. are also economically important. *Calicophoron daubneyi* belongs to the family Paramphistomidae, a group of flukes typically found in the forestomachs of ruminants. Unlike the flattened morphology common to most trematodes, these flukes exhibit a distinct conical shape as adults [12]. *C. daubneyi* is considered an emerging threat in Europe due to its impact on livestock productivity. The larval stage of rumen fluke are released into the duodenum, where they attach to the intestinal lining and causes tissue damage. Although chronic *C. daubneyi* infection is not typically associated with clinical disease, some negative effects on production have been reported [12]. Recent studies suggest that paramphistomosis is now more prevalent than fasciolosis in certain regions of the UK [9–11]. However, currently, diagnostic options for rumen fluke are limited and need further research.

Traditionally, diagnosis of both *Fasciola* and *Calicophoron* infections relies on microscopic identification of fluke eggs in the host faeces [13–15], with eggs usually observed 10−12 weeks post-infection and thereafter [14–16]. The effectiveness of microscopy relies on personnel training and expertise. Moreover, it becomes labour-intensive when handling a large number of samples, particularly if the person lacks sufficient experience, leading to low sensitivity [17]. Distinguishing between *F. hepatica* and *C. daubneyi* solely based on egg morphology in faecal samples can be challenging as both parasites produce eggs with comparable sizes and shapes [18,19]. Although, it is reported that *F. hepatica* eggs can be identified by their operculum [20] and yellowish colour [21], these features can be difficult to observe consistently using standard light microscopy. As a result, trusting solely in egg morphology for diagnosis may lead to misidentification. Molecular diagnostics based on *Fasciola* DNA detection are rapidly progressing; for instance, qPCR [22] and PCR techniques have been applied to adult worms and infected snails [23]. Additionally, techniques such as nested PCR [14] and Loop-Mediated Isothermal Amplification (LAMP) are being explored [24] and are under assessment for their speed, reliability, and accuracy compared to other methods [25]. However, significant challenges remain in detecting *Fasciola* DNA in faecal material, highlighting the need for a reliable, time-efficient and accurate diagnostic method, which can handle medium to large sample sizes and is capable of differentiating *F. hepatica* infections from other fluke species.

Next-generation sequencing technologies are transforming the diagnosis of infectious diseases whilst also paving the way for new research areas, such as microbiome studies [26–28]. Amplicon sequencing using next-generation approaches has been applied to identify gastrointestinal nematode species in ruminants (“nemabiome”) [29], to quantify trypanosome and piroplasm species in ruminant blood samples (“haemoprotobiome”) [30,31]. Similar “tremabiome” technology has been applied to quantify single species adult fluke infections in ruminants [32–34]. Despite the prevalence of mixed fluke infections in ruminants [32,33], this approach has not yet been used to understand fluke egg communities in single and mixed species infections.

This study aimed to develop a user-friendly real time diagnostic tool to detect *F. hepatica* and *F. gigantica* infections, and to distinguish them from other species such as rumen fluke. Specifically, a SYBR Green qPCR-based assay was utilised to amplify the mitochondrial NADH1 (mt-ND1) DNA marker without the need for fluorescent probes. This assay was designed to screen *Fasciola* spp. infections from faecal sedimented material of naturally infected cattle and sheep. Additionally, we combined two previously published deep amplicon sequencing approaches tested on adult flukes of *Fasciola* and *Calicophoron* spp. [32,34] to create a tremabiome approach capable of differentiating between various species of fluke. Finally, both methods were validated using field samples of fluke eggs and adults collected from different regions across the UK and compared to microscopy.

## Materials and Methods

### Ethical statement

Non-invasive collection of faecal samples was approved by the NASPA (Non-Animal Scientific Procedures Act) sub-committee of AWERB, University of Surrey, UK, under the reference NASPA-2122-04 for the project “Developing Novel Rapid Diagnostics for Neglected Parasitic Diseases.” Adult *F. hepatica* were collected at licenced slaughterhouses and through post-mortem examination. Completion of a University of Surrey SAGE-AR indicated that no formal ethical approval was required for adult fluke sampling.

### Positive control samples

All adult *F. hepatica* worms and sedimented eggs were collected from UK, whilst adult worms and eggs purified from adult worms of *F. gigantica*, *C. daubneyi*, *Paramphistomum epiclitum* and *Explanatum explanatum* were collected from abattoirs in Pakistan in our previous studies [32–36]. Adult worm tissue was processed following a previously described protocol [33] and DNA was extracted using the DNeasy Blood & Tissue Kit (Qiagen, USA).

DNA was then extracted from eggs of positive controls (*F. hepatica*, *F. gigantica*, *C. daubneyi*, *E. explanatum*) utilising the DNeasy PowerSoil Pro Kits (Qiagen, USA), with slight modification. Briefly these modifications included the incubation of sedimented material with the lysis buffer (CD1) for 10 minutes at 65°C, followed by 10 minutes of bead-beating in a TissueLyser LT (Qiagen, USA). The manufacturer’s protocol was then followed, with DNA being eluted in 10mM Tris buffer and stored at-80°C for downstream analysis.

### Field sample collection

To engage cattle and sheep farmers, the study was advertised by email to registered veterinary practitioners using the Royal College of Veterinary Surgeons Find a Vet site filtering for practices specialising in cattle, sheep and/or goats, and camelids. In addition, societies for sheep and cattle listed by the Department for Environment, Food and Rural Affairs (https://www.gov.uk/government/publications/lists-of-recognised-animal-breeding-organisations) in the UK were also approached via the email contact listed. Participants were sent a Royal Mail prepaid SafeBox with a sampling kit, a short questionnaire, and participant information sheet and consent form according to ethical requirements. No samples were used without the written informed consent of the farmer. A total of 402 faecal samples were collected from 19 cattle and sheep farms across various geographical regions of the UK through 10 registered veterinary practitioners from December 2022 to May 2024 (Table S1).

In addition to faecal samples, adult *F. hepatica* worms were collected from abattoirs and at post-mortem analysis from cattle (n=2) originating from West Sussex, and East Sussex, as well as from sheep (n=10) from West Sussex, Kent, Derbyshire, Renfrewshire Scotland and County Tyrone Northern Ireland. All samples were transported to the School of Veterinary Medicine at the University of Surrey, UK, and subsequently stored at-20°C for further analysis. DNA from the adult flukes was extracted as described in section “Positive control samples”.

### Morphological identification of fluke eggs

Faecal samples were initially processed using standard sedimentation methodology [37] and then to streamline the process a time-saving method Flukefinder® (Diagnostic System, USA) was also used [18,38], with slight adjustments. Both methods utilised 7-10 grams of faecal material which was combined with 50 ml of water, sieved through gauze and then passed through the Flukefinder® apparatus. The collected filtered material from both methods was then mixed with 250 ml of water in a conical beaker. After three minutes, the supernatant was removed; this step was repeated thrice for clarity. The sediment was transferred to a 50 ml centrifuge tube filled with water, and the supernatant was aspirated after 3 minutes. This process was repeated with a 15 ml centrifuge tube. Finally, the sediment was transferred to a 1.5 ml Eppendorf tube in 1 ml of PBS and was stored at 4°C for subsequent microscopy and DNA isolation.

Morphological egg examination involved inspecting 100 to 500 μl of sedimented faeces stained with 0.5% methylene blue (Pro-Lab, UK) in a counting chamber (Graticules Optics Limited, UK) under a compound microscope (Nikon, Japan) at 100× magnification with a lens containing a graticule. Samples positive for fluke eggs (n=128) were assessed for the presence of *F. hepatica* or *C. daubneyi* eggs based on their morphological characteristics, including size, shape, colour and operculum [18–21]. The overall workflow is summarised in Fig. 1 (a).

**Fig. 1:**
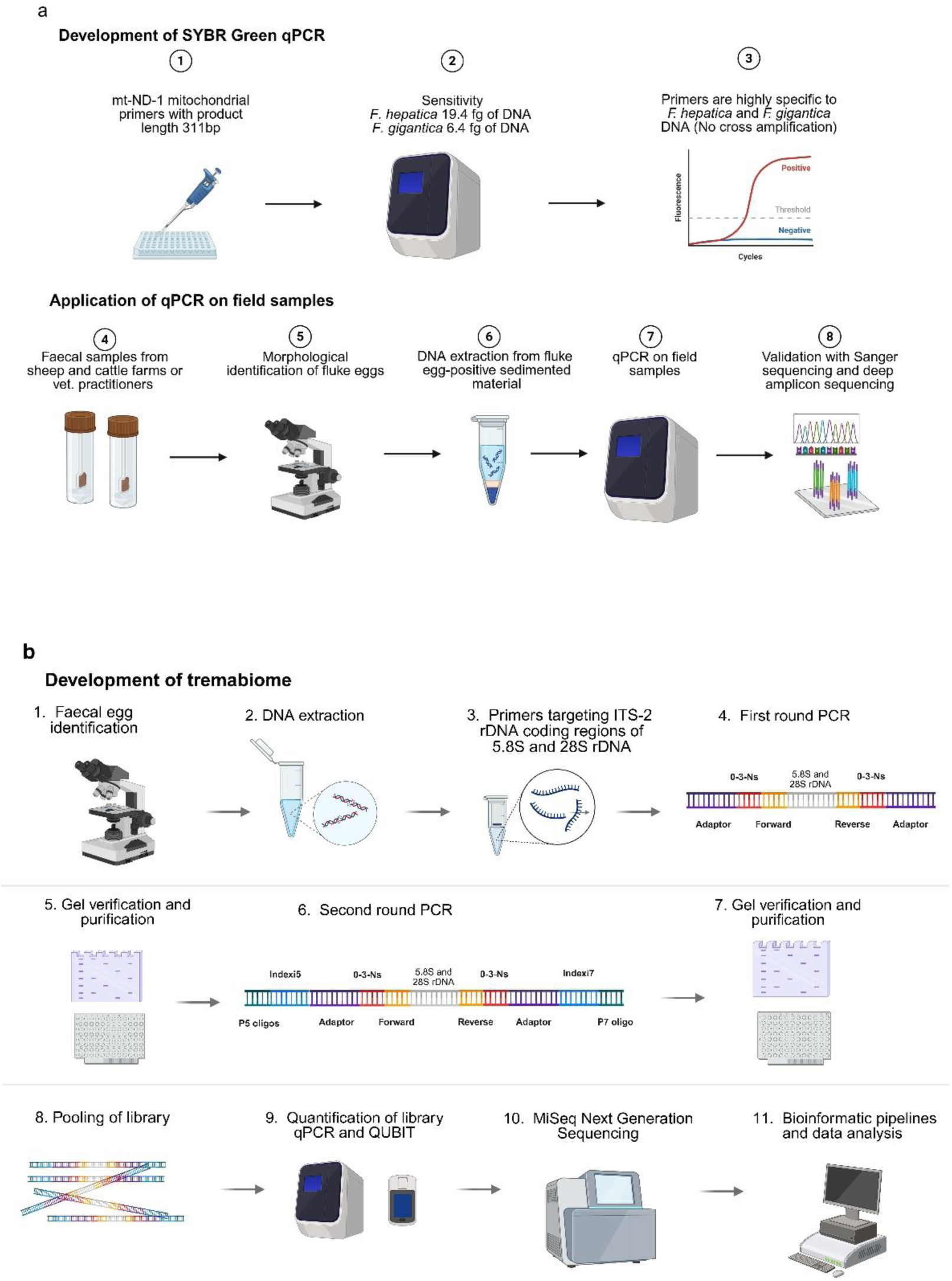
Developmemt of qPCR and tremabiome (a) Workflow adopted for developing qPCR and screening of faecal samples for the presence of *F. hepatica* infection. (b) Overview of development of the tremabiome method and its application on natural fluke infections in ruminants in the UK.

Any sedimented faecal samples which were identified as positive for fluke eggs by microscopy were subsequently subjected to DNA isolation utilising the DNeasy PowerSoil Pro Kits (Qiagen, USA) according to the manufacturer’s instructions as described in the section “Positive control samples”.

### Molecular identification of fluke eggs

PCR was performed using universal ITS2 primers [39] on selected fluke egg-positive samples and all positive controls. Further, *Fasciola* specific mt-ND1 primers [33] were applied to selected samples to confirm the diagnostic accuracy of the PCR targets. This was followed by Sanger sequencing to confirm the presence and correct amplification of DNA for different fluke species.

All PCR reactions were prepared with DreamTaq Green PCR master mix (Thermo Scientific, USA) in a 25 μl reaction mix, with primer concentrations of 200 nM and 4 μl of sample DNA template. The PCR cycling conditions were initial denaturation at 95°C for 5 minutes, followed by 35 cycles of denaturation at 95°C for 1 minute, annealing at 55°C (ITS2 primers), 50°C (mt-ND1 primers) for 1 minute, and extension at 72°C for 1 minute. The final extension step was carried out at 72°C for 5 minutes. Positive controls consisted of DNA from *F. hepatica* and *F. gigantica* adult worms. The resulting PCR products were purified and cleaned using a NucleoMag kit for clean-up and size selection of NGS library prep reactions (MACHEREY-NAGEL, GmbH & Co.KG). Sanger sequencing of the PCR product was performed by Source Biosciences, UK and Eurofins Genomics, Germany. Selected samples were subjected to conventional PCR and Sanger sequencing due to limited resources. All obtained sequences were visualised using Geneious version 8.0.5 (https://www.geneious.com), and the FASTA sequences were submited to the BLASTn tool on NCBI to confirm fluke species identity.

### Development and validation of a SYBR green qPCR to detect *Fasciola* eggs at species level

A SYBR green qPCR assay was developed to detect *Fasciola* spp, ingus mt-ND1 primers. These mt-ND1 primers were previously designed and employed in a meta-barcoded PCR [33]. In the SYBR green assay, 3 μl of DNA was subjected to qPCR using 10 μl of 2X SsoAdvanced Universal SYBR Green Supermix (Bio-Rad, USA), resulting in a total reaction mix volume of 20 μL, including 500 nM of mt-ND1 primers.

The cycling program was initial denaturation at 98 ℃ for 3 mins, followed by 40 cycles of denaturation at 98 ℃ for 15 secs and annealing at 60 ℃ for 30 secs on a CFX96 Real-Time PCR machine (Bio-Rad, USA). Subsequently, a melt curve was generated from 65 ℃ to 95 ℃ with an increment of 0.5 ℃ for 0.05 secs per plate read. Positive and negative controls were employed as described above. All samples were subjected to qPCR in triplicate, and the resulting data were visualised using CFX Maestro Version: 5.3.022.1030 (Bio-Rad, USA).

Detection sensitivity limits of the assay were assessed using five-fold serial dilutions of *F. hepatica* (ranging from 300 pg to 0.768 fg) and *F. gigantica* (ranging from 500 pg to 1.28 fg) adult worm DNA. The DNA was quantified using Qubit™ dsDNA HS and BR Assay Kits (Invitrogen™). The analytical specificity of the assays was evaluated by testing 1 ng of DNA from other prevalent flukes and nematodes found in sheep and cattle. These included *C. daubneyi*, *P. epiclitum*, *E. explanatum*, and the nematode *Teladorsagia circumcincta.* The sensitivity and specificity tests were conducted in triplicate and repeated twice.

The reliability of the method was measured by comparing inter-and intra-assay variations in Cq values for *F. hepatica* and *F. gigantica* DNA. For validation, the newly developed qPCR assay was applied to egg DNA extracted from sedimented faecal samples.

### Development and validation of deep amplicon sequencing to detect fluke eggs at species level

A universal ITS2 rDNA marker for a range of 21 different fluke species (Table S2; DOI: 10.17632/zyvwc6ppy8.1) targeting coding regions of 5.8S and 28S rDNA was used, expected to produce a fragment of 490–743 bp [39]. The primers were meta-barcoded by adding Illumina adaptor sequences to the both forward and reverse primers, along with up to three random’N’ nucleotides positioned between the adaptor sequences and the locus-specific primers. Additionally, modified phosphate bonds were added between the last three nucleotides of each primer to enhance their stability (Table S3) and used in PCR amplification to detect fluke species. To assess the representation of species read depth, mock DNA mixtures were prepared in triplicate by pooling 250 eggs of each species, including *F. hepatica*, *F. gigantica*, and *C. daubneyi* and subjected to amplicon sequencing in triplicate. PCR cycle numbers were adjusted 35x, 30x and 25x in the first round to examine their effect on species representation. Further, to evaluate species representation using egg DNA, we created seven mock egg pools in triplicate, adjusting the proportion of *F. hepatica* and *C. daubneyi* in an approximate total of 250 eggs with ratios of 99:1, 90:10, 70:30, 50:50, 30:70, 10:90, and 1:99. Moreover, to test the threshold of deep amplicon sequencing, pools of equal egg proportions (50 eggs) were prepared from three out of four species (*F. hepatica*, *F. gigantica*, *E. explanatum*, *C. daubneyi*). The fourth species (*F. hepatica* or *C. daubneyi*) was then added in decreasing numbers of 500, 50, 20, 15, 5, and 0 eggs, creating six mock pools. The first round of PCR was performed using the KAPA HiFi PCR Kit (KAPA BIOSYSTEMS, South Africa). The modified primer sets, adaptors, barcoded PCR amplification conditions, magnetic bead purification methods and bioinformatic analysis were based on our previously described methods [33]. The first-round PCR products were subjected to a second-round PCR using a barcoded primer set to attach a unique barcode index fragment required for Illumina sequencing [30] Fig. 1 (b).

10 μl of each second round barcoded PCR product were combined to create a pooled library and then purified by agarose gel electrophoresis to remove non-specific products and adaptor dimers. During post-run processing, the MiSeq system separated all sequencing data by sample quality using the barcoded indices to generate FASTQ files (raw sequence read files available at Mendeley database DOI: 10.17632/zyvwc6ppy8.1), see workflow diagram Fig. 1 (b).

The FASTQ files obtained from the post-run Illumina MiSeq (BioProject ID PRJNA1273189) were analysed in Mothur/1.41.0-Python-2.7.15 [40] using the High-Performance Cluster (HPC) system at the University of Surrey, UK. Pipelines described in our previous study [41] were utilised with modifications of the newly developed reference sequence library (Script available at DOI: 10.17632/zyvwc6ppy8.1).

To generate a taxonomy file, ITS2 rDNA reference sequences (n=545) were obtained from NCBI, representing 21 fluke species (Table S2). The genetic distances between different fluke species were then calculated based on the sequenced region. Overall, a variation in genetic identity was found between different fluke species ranging from 40% to 99% (Table S4, DOI: 10.17632/zyvwc6ppy8.1). Finally, a phylogenetic tree of 154 unique sequences showed a distinct clustering of each fluke species (Fig. 2) as described in the section phylogenetic analysis.

**Fig. 2:**
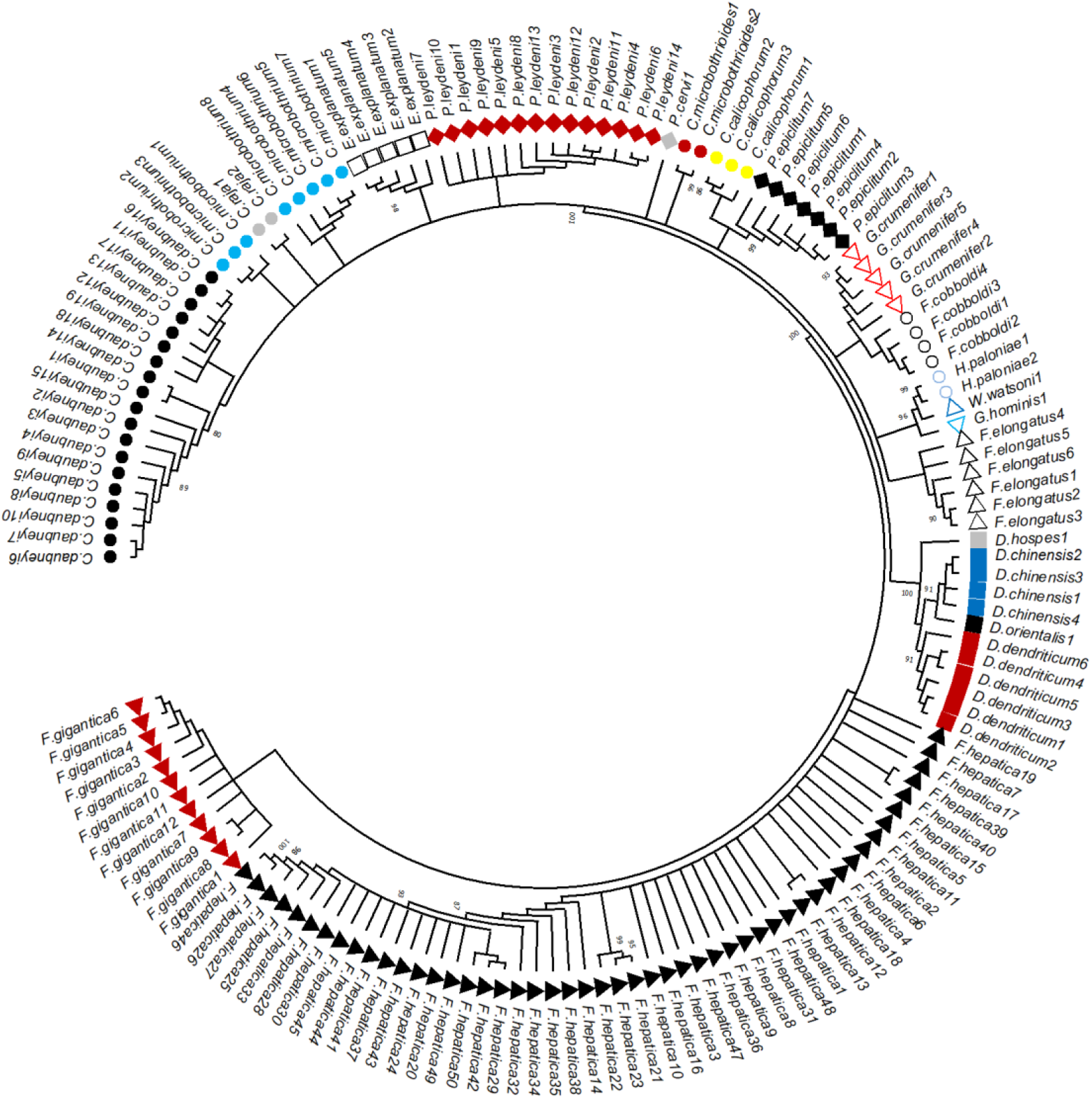
Neighbor-Joining tree generated for fluke species including reference sequences of 21 different fluke species. The reference sequences were downloaded from NCBI data, and 154 unique sequences were selected and aligned (Table S2, DOI: 10.17632/zyvwc6ppy8.1). Different fluke species are indicated with symbols of different colours and shapes. *F. hepatica* red triangle, *F. gigantica* black triangle, *C. daubneyi* black circle, *Paramphistomum leydeni* red diamond, *P. cervi* grey diamond, *C. microbothrium* light blue circle, *P. epiclitum* black diamond, *Gastrothylax crumenifer* triangle with red boundary no fill, *Fischoederius elongatus* triangle with black boundary no fill, *Dicrocoelium dendriticum* red square, *C. calicophorum* yellow circle, *Fischoederius cobboldi* circle with black boundary no fill, *C. microbothrioides* red circle, *Homalogaster paloniae* circle with light blue boundary no fill, *E. explanatum* square with black boundary no fill, *C. raja* grey circle, Watsonius watsoni triangle with dark blue border no fill, *Gastrodiscoides hominis* triangle with light blue boundary no fill, *D. orientalis* black square, *D. hospes* grey square, and *Dicrocoelium chinensis* dark blue square.

Following extraction of the taxonomy file, quality filtering was conducted for the identification of unique fluke sequences, detailed count tables were generated and an alignment (ALIGN) file of sequences across all samples was produced. This workflow ensured a robust, high-quality dataset suitable for downstream taxonomic studies for flukes using a series of commands (Script available at DOI: 10.17632/zyvwc6ppy8.1).

The count table and alignment (ALIGN) file were further used for the extraction of amplicon sequence variants (ASVs). The R script extracted specific ASVs corresponding to the target species of flukes applying cutoff values of 250 reads each (NCBI accession numbers; PV752375-PV752429, PV752431-PV752462). The R script started by cleaning sequence names, trimming whitespace and converting them to lowercase to ensure consistency for successful merging. After identifying unmatched sequences, the datasets were merged based on sequence names. The merged data was filtered to retain only rows with a ‘total’ count of at least 250 reads, ensuring that only high-quality consensus sequences remained. A custom function was employed to write the sequences into a combined FASTA file, preserving both the sequence names and read counts. Next, the script utilised the Biostrings package to clean the sequences by removing ‘N’ characters and ambiguous bases, saving the high-quality sequences to a final FASTA file (R script available at DOI: 10.17632/zyvwc6ppy8.1). This comprehensive approach ensured accurate sequence extraction for subsequent analysis. Finally, the sample-wise separated FASTA files were subjected to remote NCBI BLASTn loop command using “blastn: 2.16.0+” in the HPC cluster system (command line available DOI: 10.17632/zyvwc6ppy8.1). Mismatched sequences with the NCBI database were considered contaminated sequences and discarded.

### Statistical analysis of qPCR and deep amplicon sequencing data

For qPCR analysis, the raw Cq values were extracted from CFX Maestro Version: 5.3.022.1030 (Bio-Rad, USA) and a linear standard curve was created by plotting DNA quantities against the average Cq values for each concentration tested. For deep amplicon sequence reads data analysis, the sequences from the bioinformatics pipeline were further analysed for sequence accuracy and percentage identity using remote blastn: 2.16.0+ with the NCBI database. To determine the percentage composition of *F. hepatica*, *F. gigantica*, *C. daubneyi*, and *E. explanatum* in mock egg mixtures (positive control) and field samples, species composition percentages were calculated by dividing the classified sequence reads for each species by the total reads per sample. A one-way ANOVA was applied to assess the proportional representation of mock mixes comprising different fluke species eggs across varying PCR amplification cycles. The association between qPCR and tremabiome results was assessed using Chi-squared analysis. All visualisations of data were performed in R version 4.3.3 (R scripts are available at DOI: 10.17632/zyvwc6ppy8.1)

### Phylogenetic analysis

Phylogenetic trees were generated from unique reference sequences of ITS2 rDNA from 21 different fluke species downloaded from NCBI GenBank (Table S2). The sequences were aligned using the MUSCLE alignment tool in Geneious v8.0.5 (Biomatters Ltd, New Zealand) and genetic distances were calculated (Table S4). Further, a phylogenetic tree of the unique ITS2 rDNA sequences for all 21 fluke species and ASVs of the flukes was constructed using the Neighbor-Joining method [42]. The evolutionary distances were computed using the Maximum Composite Likelihood method [43] in MEGA11 [44] with a bootstrap value of 2000 [45].

## Results

### Fluke identification by microscopy

A total of 402 faecal samples were examined, out of which 128 were positive for fluke eggs. The sampled animals included cattle (n=154), sheep (n=233), water buffalo (n=1), alpaca (n=4), and goats (n=2), with animal species unspecified for 8 samples. Of these samples, 191 had a history of *Fasciola* infection, 119 had no history, and 92 had an unknown history (Table S1). The cattle and sheep sampled represented diverse age groups, ranging from calves and lambs to adults, providing a comprehensive representation of the animal host population. Notably, only one sheep farm from Wales was included, and no faecal samples were collected from Northern Ireland. Based on egg morphology and staining, 30 samples were identified as *C. daubneyi*, 71 as *F. hepatica*, five as mixed infection, and 22 remained undecided (Table S5).

### PCR and Sanger sequencing for fluke identification and comparison with microscopy

Out of 128 samples positive for fluke eggs, DNA was successfully extracted from 125. PCR was performed on 85 randomly selected samples using universal ITS2 primers, with bands observed in 73 samples. From the 71 samples believed to be *F. hepatica* by microscopy, 37 were screened by PCR, of which 31 samples were analysed by Sanger sequencing. Of these 31, three were confirmed as *F. hepatica*, 19 as *C. daubneyi* and nine samples demonstrated poor sequence quality. Of the 30 *C. daubneyi* positive samples identified by microscopy, 30 were screened by PCR, of which 27 samples were then analysed by Sanger Sequencing. Of these 27 samples, 16 were confirmed as *C. daubneyi*, one was confirmed to be *F. hepatica*, one was identified as *Paramphistomum epiclitum*, and nine demonstrated poor sequence quality. Of the five samples which were identified as mixed infections by microscopy (*F. hepatica* and *C. daubneyi*), three were screened by PCR and analysed by Sanger sequencing. Of these three mixed infection samples, Sanger sequencing identified one sample as *F. hepatica*, and two as *C. daubneyi*. In the 22 samples where microscopy could not determine the fluke species present, 15 samples were screened by PCR, of which 14 were subsequently analysed by Sanger sequencing. From these, two samples were confirmed as *F. hepatica*, six as *C. daubneyi*, and six demonstrated poor sequence quality (Table 1, Table S5).

**Table 1:** Comparison of microscopy, ITS2 PCR and Sanger sequencing on fluke egg-positive samples.

| Microscopy | PCR |  |  |  |  | Sanger sequencing using ITS2 |  |  |  |  |  |  |
| --- | --- | --- | --- | --- | --- | --- | --- | --- | --- | --- | --- | --- |
|  | Negative | Positive | Not performed | ND<sup>2</sup> | Total | *F. hepatica* | *C. daubneyi* | *Paramphistomum epiclitum* | PSQ | Not performed | ND<sup>2</sup> | Total |
| *F. hepatica* | 7 | 30 | 32 | 2 | 71 | 3 | 19 | 0 | 9 | 38 | 2 | 71 |
| *C. daubneyi* | 4 | 26 | 0 | 0 | 30 | 1 | 16 | 1 | 9 | 3 | 0 | 30 |
| Mixed | 0 | 3 | 2 | 0 | 5 | 1 | 2 | 0 | 0 | 2 | 0 | 5 |
| Fluke<sup>1</sup> | 1 | 14 | 6 | 1 | 22 | 2 | 6 | 0 | 6 | 7 | 1 | 22 |
| Total | 12 | 73 | 40 | 3 | 128 | 7 | 43 | 1 | 24 | 50 | 3 | 128 |

<sup>1</sup>Unidentified flukes; <sup>2</sup> ND = DNA extraction failed; PSQ = poor sequence quality

In addition to the ITS2 PCR and Sanger sequencing, a third PCR based diagnostic test, targeting mitochondrial target mt-ND1 and specific for identifying *Fasciola* spp. was utilised. For this, 15 samples were randomly selected within the diagnostic categories already created (*F. hepatica*, *C. daubneyi*, mixed and unknown). From the two morphologically identified *C. daubneyi* samples selected, one was confirmed as *F. hepatica*, while the other had poor sequence quality. For instance, from the nine morphologically identified *F. hepatica* samples, two were initially identified as *F. hepatica* but were confirmed as *C. daubneyi* by ITS2, whereas mt-ND1 sequencing confirmed them as *F. hepatica*. This suggests that microscopy failed to identify mixed infections in those two samples. Additionally, one sample had poor ITS2 sequence quality but was confirmed as *F. hepatica* by mt-ND1 and three samples were not sequenced for ITS2 but were confirmed as *F. hepatica* by mt-ND1.

Additionally, one morphologically identified mixed infection by microscopy was supported by ITS2 and ND1 PCR assays and subsequent Sanger sequencing, confirming the presence of *C. daubneyi* and *F. hepatica*, respectively. Among the three morphologically unidentified samples, one was confirmed as *C. daubneyi* by ITS2, with mt-ND1 sequencing demonstrating poor sequence quality, while another was not sequenced for ITS2 but confirmed as *F. hepatica* by mt-ND1. (Table 2, Table S5). These findings highlight discrepancies between morphological identification and molecular confirmation using a nuclear and mitochondrial DNA target, emphasising the need for more accurate molecular techniques.

**Table 2:** Comparison of microscopy and Sanger sequencing using mt-ND-1 markers on fluke egg-positive samples.

| Microscopy | Sanger sequencing using ND1 |  |  |  |  |  |
| --- | --- | --- | --- | --- | --- | --- |
|  | *F. hepatica* | *C. daubneyi* | PSQ | Not performed | ND<sup>2</sup> | Total |
| *F. hepatica* | 6 | 0 | 3 | 60 | 2 | 71 |
| *C. daubneyi* | 1 | 0 | 1 | 28 | 0 | 30 |
| Mixed | 1 | 0 | 0 | 4 | 0 | 5 |
| Fluke<sup>1</sup> | 1 | 0 | 2 | 18 | 1 | 22 |
| Total | 9 | 0 | 6 | 110 | 3 | 128 |

<sup>1</sup>Unidentified flukes; <sup>2</sup> ND = DNA extraction failed; PSQ = poor sequence quality

### Detection of *Fasciola* species using a newly developed qPCR assay

To provide a simple, low-cost, sensitive, universal, and accurate molecular method for diagnosing *Fasciola* infections in faeces, a SYBR green qPCR assay was developed using a repurposed primer set targeting mt-ND1 specific to *F. hepatica* and *F. gigantica.* The assay’s analytical sensitivity was first assessed using quantified *F. hepatica* and *F. gigantica* adult worm DNA (positive controls). The assay’s limit of detection based on a 1:5 DNA dilution series was found to be 19.2 fg for *F. hepatica* and 6.4 fg for *F. gigantica* DNA. A linear standard curve was generated, showing an efficiency of 97% (R² = 0.9759) for *F. hepatica* and 99% (R² = 0.9995) for *F. gigantica,* demonstrating efficient primer binding and target amplification. Additionally, qPCR melt curve analysis identified distinct peaks at 81.50°C for *F. hepatica* and *F. gigantica* (Fig. S1, A and B), confirming the specificity of primer binding to the same DNA target, and absence of nonspecific primer interactions. The specificity of the qPCR assay was evaluated against DNA from other prevalent fluke and nematode species. The melt curve analysis confirmed that only *F. hepatica* and *F. gigantica* produced amplification peaks at 81.50°C, with no cross-amplification (Fig. S2). The assay exhibited strong reproducibility, with coefficients of variation (CV) below 6.0% for intra-assay and inter-assay. The mean Cq values ranged from 21.96 to 38.26 for *F.* hepatica with DNA dilutions ranging from 300 pg to 19.2 fg and 18.62 to 38.24 for *F. gigantica* with DNA dilutions ranging from 500 pg to 6.4 fg), maintaining consistency across replicates (Table S6).

To gain further clarification on which *Fasciola* species were present in the 128 egg-positive field samples obtained, the newly developed specific SYBR green qPCR was utilised. Of the 128 samples screened, 57 were positive for *F. hepatica*, 66 were negative, and five were either not determined or failed DNA extraction.

### Comparison of microscopy and qPCR for fluke species diagnosis

Among the 71 samples identified as *F. hepatica* positive by microscopy, qPCR confirmed *F. hepatica* in 34 samples, while 33 samples were negative, and four were not determined or failed DNA extraction. From the 30 samples identified as *C. daubneyi* by microscopy, qPCR detected *F. hepatica* in seven samples, whereas 23 samples tested negative. From the five samples identified as mixed infections by microscopy, qPCR confirmed four as positive for *F. hepatica* infections, while one sample tested negative. Finally, of the 22 samples which were undecided using microscopy, qPCR identified 12 as *F. hepatica*, nine as negative, and one sample was not determined or with failed DNA extraction (Table 3, Table S5).

**Table 3:** Comparison of microscopy, qPCR (for *F. hepatica*) and tremabiome on 128 fluke egg-positive samples.

| Microscopy | qPCR |  |  |  |  | Tremabiome |  |  |  |  |  |
| --- | --- | --- | --- | --- | --- | --- | --- | --- | --- | --- | --- |
|  | Negative | Positive | ND | NP | Total | *F. hepatica* | *C. daubneyi* | Mixed | ND<sup>2</sup> | No reads | Total |
| *F. hepatica* | 33 | 34 | 2 | 2 | 71 | 11 | 26 | 30 | 2 | 2 | 71 |
| *C. daubneyi* | 23 | 7 | 0 | 0 | 30 | 2 | 21 | 7 | 0 | 0 | 30 |
| Mixed | 1 | 4 | 0 | 0 | 5 | 0 | 1 | 4 | 0 | 0 | 5 |
| Fluke<sup>1</sup> | 9 | 12 | 1 | 0 | 22 | 0 | 3 | 17 | 1 | 1 | 22 |
| Total | 66 | 57 | 3 | 2 | 128 | 13 | 51 | 58 | 3 | 3 | 128 |

<sup>1</sup>Unidentified flukes; <sup>2</sup> ND = DNA extraction failed; NP=Not performed

### Detection of fluke species using a newly developed tremabiome deep amplicon sequencing method

To accurately identify mixed-species as well as single-species fluke infections, a tremabiome deep amplicon sequencing assay was developed using a universal primer set targeting rDNA ITS2, specific to fluke species. Initially, the sequence representation of three different fluke species *F. hepatica*, *F. gigantica*, and *C. daubneyi* in the deep amplicon sequencing assay was determined (Fig. 3a). This allowed the evaluation of proportional DNA sequence output reads relative to known species ratios. Each species showed significant representation in the sequence counts in each mix (Fig. 3a). Furthermore, the number of cycles (25X, 30X and 35X) used during the adaptor PCR were validated to ensure sufficient DNA was generated for sequencing whilst maintaining a balance between amplification efficiency and accuracy. Whilst it is known that an appropriate number of cycles helps minimise deviations and PCR bias, and over-amplification causing sequence dominance, we found that the number of cycles did not affect the sequence representation of any species. In each mock pool, *C. daubneyi* generated the highest number of sequence reads, followed by *F. gigantica* and *F. hepatica*. Despite these trends, no statistically significant differences were observed in the proportional representation of any species across the different PCR amplification cycles (*F. gigantica*, *P* = 0.730; *F. hepatica*, *P* = 0.774; and *C. daubneyi*, *P* = 0.258) (Fig. 3a).

**Fig. 3:**
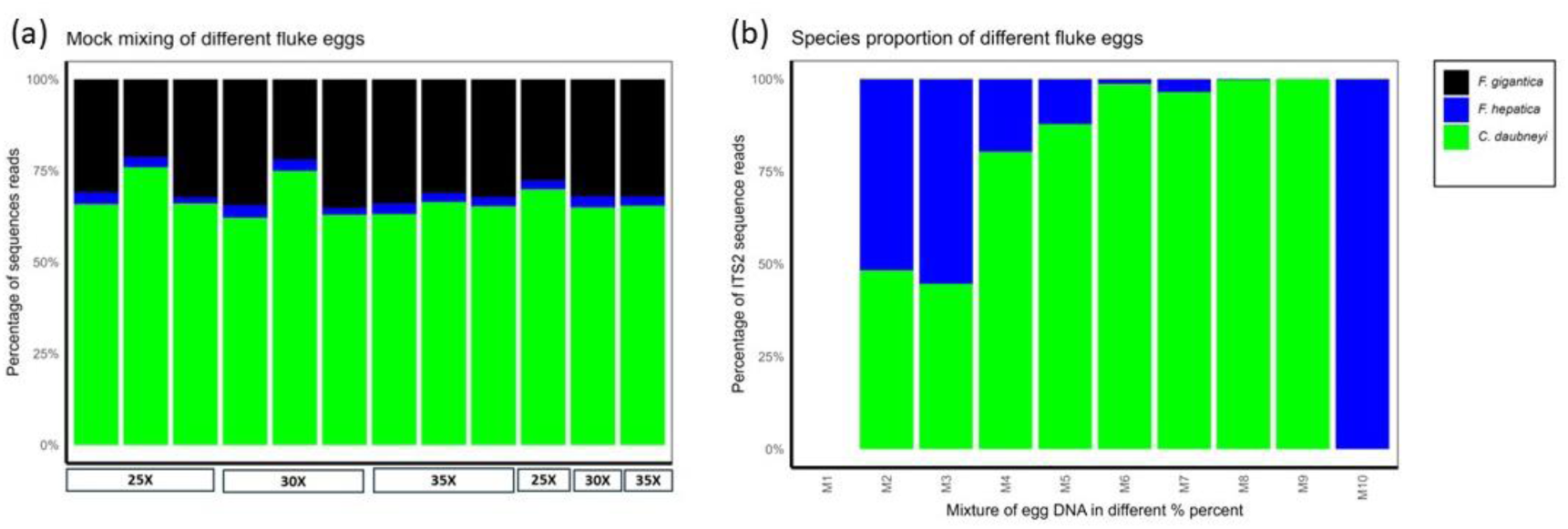
Sequence representation of the mock mixture of fluke species in deep amplicon sequencing. (a) *F. hepatica*, *F. gigantica*, and *C. daubneyi*. DNA was extracted in triplicate from pooled samples containing 250 eggs of each species. The DNA mixture was amplified using PCR at three cycle levels (25X, 30X, and 35X), with triplicate testing for each pool. The x-axis indicates PCR cycle numbers, while the y-axis represents each species’ percentage of ITS2 rDNA sequence reads. Triplicates were averaged and grouped based on the amplification cycles in the last three columns. (b) Relative proportions of *F. hepatica* and *C. daubneyi* in egg DNA mixtures were assessed using deep amplicon sequencing. DNA was extracted from mock pools containing varying ratios of these two fluke species, enabling evaluation of the assay’s accuracy across a range of species proportions. The x-axis represents egg mixtures with varying *F. hepatica*: *C. daubneyi* ratios: M1 (negative control), M2 (99:1), M3 (90:10), M4 (70:30), M5 (50:50), M6 (30:70), M7 (10:90), M8 (1:99), M9 (100% *C. daubneyi*), and M10 (100% *F. hepatica*). The y-axis shows the percentage of ITS2 rDNA sequence reads for each species.

We further assessed the assay’s accuracy in identifying relative species proportions in mixed infections by testing pairwise mixtures of *F. hepatica* and *C. daubneyi* eggs (Fig. 3b). This range of mixes enabled thorough validation of the sequencing assay, demonstrating reliable detection of the two species across egg ratios, which is essential for accurately identifying species representation in mixed infections. These species were selected due to their high prevalence, frequent co-occurrence in UK cattle and sheep herds, and availability in our laboratory (Fig. 3b, File S1). This approach addressed the sensitivity of deep amplicon sequencing assays in detecting trace-level amplicons. We observed minimal variation in species representation across different mixes, which did not affect the overall interpretation of relative species abundance. For example, the 99% *F. hepatica:* 1% *C. daubneyi* mix and the 90% *F. hepatica:* 10% *C. daubneyi* mix displayed similar species representationsfurther. We tested the thresholds of the deep amplicon assay for fluke egg DNA with decreasing egg levels in mixed populations, for example, *F. hepatica* and *C. daubneyi* (Fig. 4a and 4b, File S2, and File S3). A notable observation was the production of a lower number of sequenced reads, particularly for *F. hepatica*; however, this does not affect the identification. Importantly, the assay detected both *F. hepatica* and *C. daubneyi* DNA at levels down to 5 eggs per pool.

**Fig. 4.**
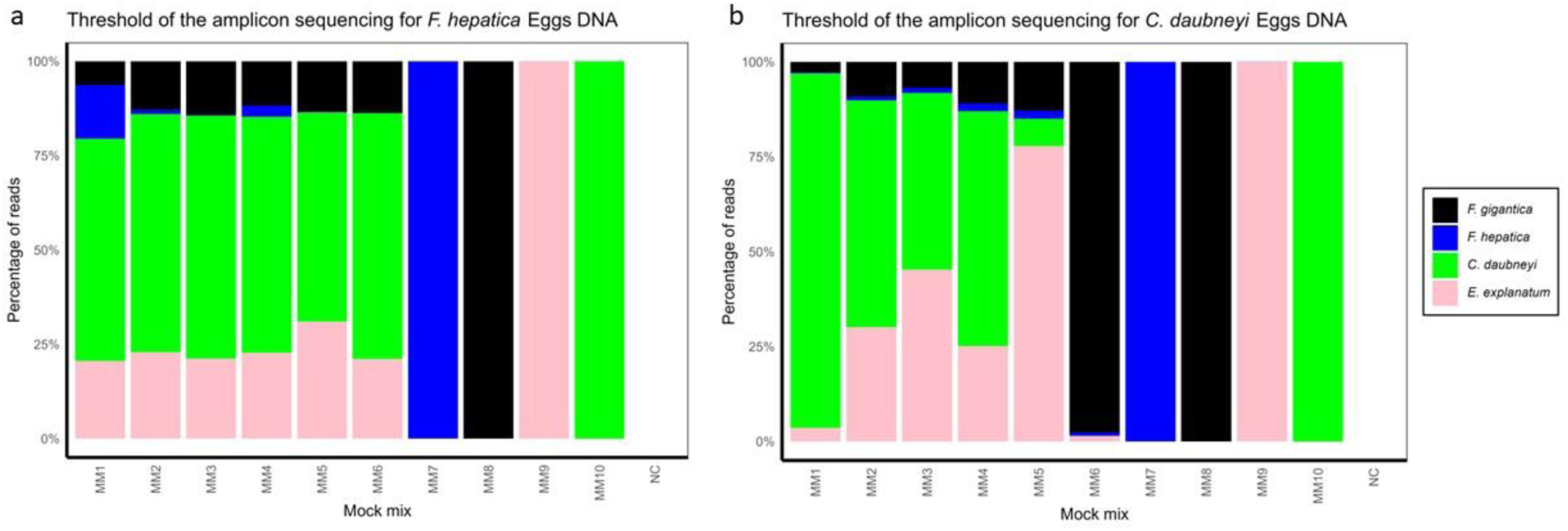
Threshold of deep amplicon sequencing Application of deep amplicon sequencing to mock egg mixtures with gradually lower counts of *F. hepatica* and *C. daubneyi* eggs. Two sets of mixtures were designed using eggs from four fluke species (*F. hepatica*, *F. gigantica*, *E. explanatum*, and *C. daubneyi*). In panel (a), which focuses on *F. hepatica*, MM1 contained 500 eggs of *F. hepatica* along with 50 eggs of each of the other three species, creating a high relative abundance of *F. hepatica*. In mixtures MM2 through MM6, the number of *F. hepatica* eggs was reduced to 50, 20, 15 5, and 0 eggs, respectively, while the counts for the other three fluke species remained constant at 50 eggs. Panel (b) follows a similar design but targets *C. daubneyi*: MM1 contained 500 eggs of *C. daubneyi* plus 50 eggs each of *F. hepatica*, *F. gigantica*, and *E. explanatum*, and in mixtures MM2 to MM6 the number of *C. daubneyi* eggs was reduced to 50, 20, 15, 5, and 0 eggs, with the other species maintained at 50 eggs each. Additionally, single-species control pools were included as MM7 (*F. hepatica*), MM8 (*F. gigantica*), MM9 (*E. explanatum*), and MM10 (*C. daubneyi*), each containing 50 eggs. The assay results show the ability to detect and accurately quantify trace levels of target DNA in mixed fluke egg populations.

After validation, the assay was applied to 125 of the 128 fluke egg-positive samples to analyse fluke species distributions in natural infections in cattle and sheep. In cattle, 67 samples (eggs: n=65, worms: n=2) produced sequence reads. The data revealed that *F. hepatica* was present in 4 samples (eggs: n=2, worms: n=2), *C. daubneyi* in 28, and mixed infections in 35 faecal samples. Similarly, in sheep, 67 samples (eggs: n=57, worms: n=10) generated sequence reads. The data showed *F. hepatica* in 21 samples (eggs: n=11, worms: n=10), *C. daubneyi* in 23, and mixed infections in 23 faecal samples (Fig. 5, File S4). Notably, out of the 125 faecal samples sequenced, 122 produced reads, as 3 samples failed to yield sequencing reads. This analysis highlights a higher prevalence of mixed infections followed by *C. daubneyi* and *F. hepatica* singular infections in cattle and sheep. The sequence reads of all samples were aligned with *F.* hepatica and *C. daubneyi* with BLASTn (DOI: 10.17632/zyvwc6ppy8.1).

**Fig. 5:**
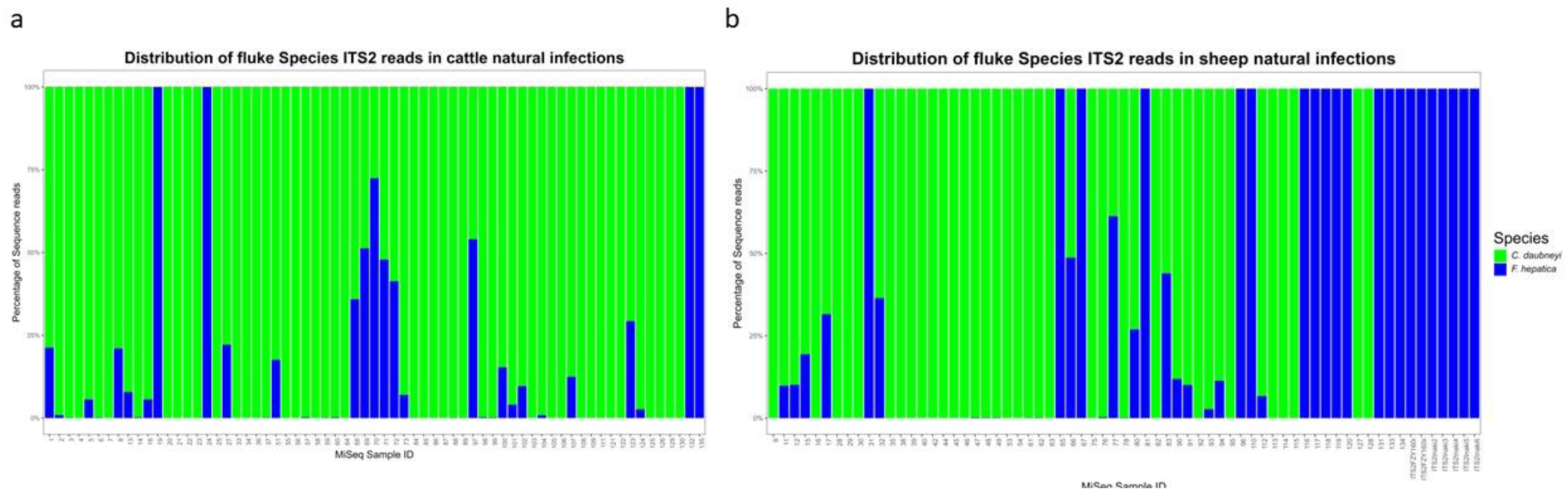
Tremabiome deep amplicon sequencing application on the field samples This figure illustrates the application of the tremabiome deep amplicon sequencing assay on DNA extracted from sedimented faecal eggs and adult worm populations collected from cattle and sheep across various regions in the UK. The charts display species proportions based on the percentage of sequence reads generated after 35 amplification cycles. Percentages of *F. hepatica* are represented in blue and *C. daubneyi* in green on the Y-axis. (a) Samples from cattle. (b) Samples from sheep.

Finally, ASVs were generated for *C. daubneyi* and *F. hepatica* from all sequence reads of the samples collected from different counties in the UK. ASVs were up to 461 bp and 527 bp for *C. daubneyi* and *F. hepatica*, respectively. In total, 87 ASVs were identified, including *F. hepatica* (n=55) and *C. daubneyi* (n=32) (DOI: 10.17632/zyvwc6ppy8.1). A phylogenetic tree of all ASVs with reference sequences of 21 fluke species (outlined in the methodology section) showed that *F. hepatica* and *C. daubneyi* species separated into distinct clades (Fig. 6).

**Fig. 6:**
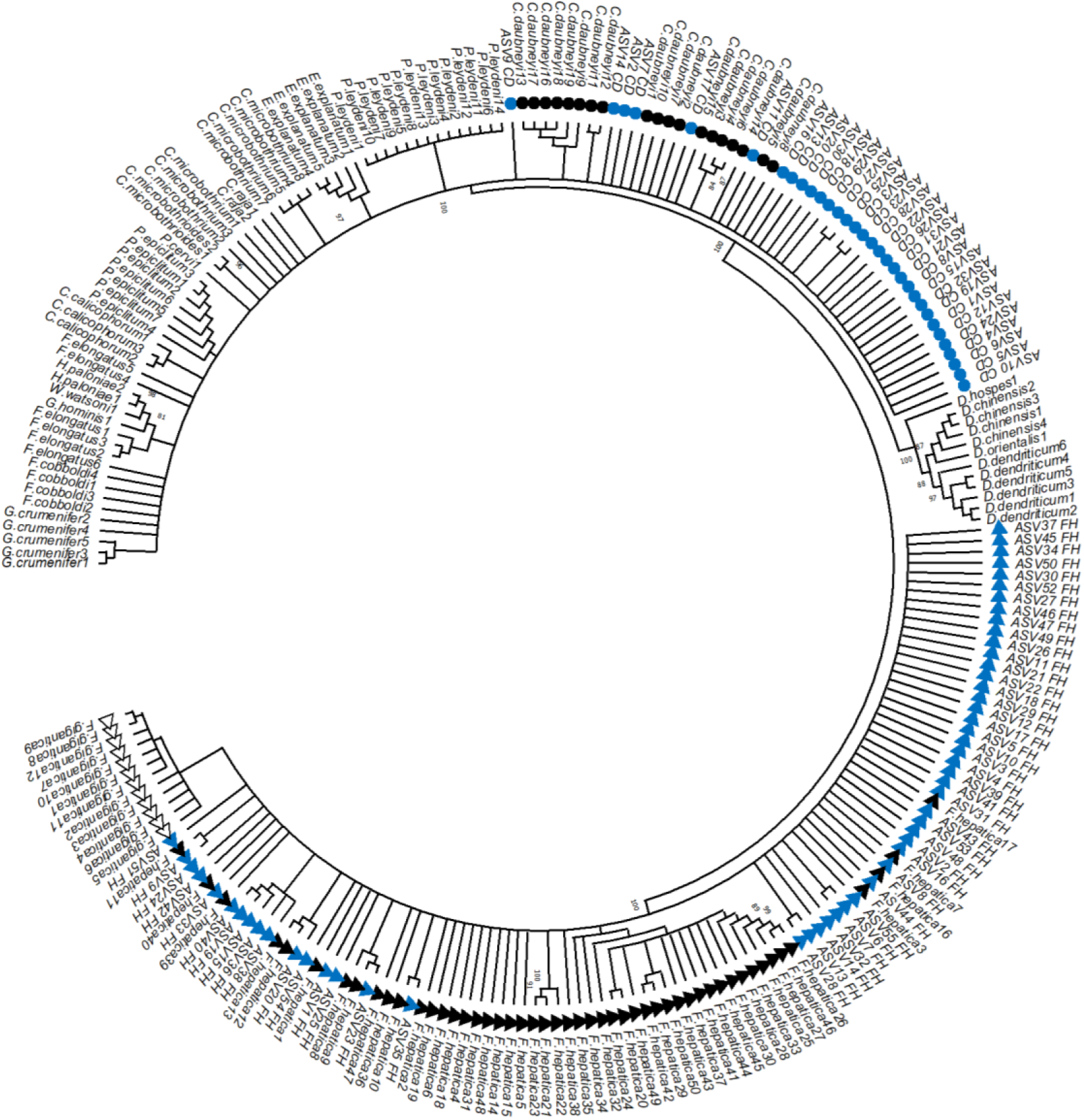
A Neighbor-Joining tree of rDNA ITS2 sequences constructed using 154 reference sequences of different fluke species downloaded from the NCBI database, along with 87 ASVs identified in this study 55 from *F. hepatica* and 32 from *C. daubneyi*. ASVs corresponding to *F. hepatica* are marked with blue triangles, while those of *C. daubneyi* are represented by blue circles. The ASVs clustered closely with their respective reference taxa, confirming accurate taxonomic assignment.

### Comparison of microscopy and tremabiome deep amplicon sequening for fluke species diagnosis

Of the 71 samples identified as *F. hepatica* by microscopy, tremabiome detected only 11 as single *F. hepatica* infections. However, tremabiome classified 26 of the 71 samples as *C. daubneyi* and 30 as mixed infections, suggesting that some samples identified as *F. hepatica* by qPCR (n=34) were mixed infections. Among the 30 samples marked as *C. daubneyi* by microscopy, tremabiome detected two as *F. hepatica*, 21 as *C. daubneyi*, and seven as mixed infections. For the microscopically recognised five mixed infections, tremabiome confirmed four as mixed infections, while one was classified as *C. daubneyi*. Lastly, among the 22 samples that were undecided by microscopy, tremabiome identified three as *C. daubneyi*, 17 as mixed infections, and two as not determined or with failed DNA extraction (Table. 3, Table S5).

### Comparison of qPCR and tremabiome deep amplicon sequencing for fluke species diagnosis

A significant correlation (*p*<0.001) was noted between the identification of *F. hepatica* infections using qPCR and the tremabiome approach. Nevertheless, there were 20 samples which tested negative by qPCR, but *F. hepatica* sequences were detected by deep amplicon (three single and 17 mixed infections). Conversely, deep amplicon sequencing did not produce *F. hepatica* reads for 8 qPCR-positive samples (Table 4, Table S5). For these samples, FECs ranged from 1-221 eggs per gram of faecal material, but due to mixed infections of *F. hepatica* and *C. daubneyi*, the egg counts for individual fluke species are not clear. Interestingly, *C. daubneyi* reads were generated for the 8 samples, which were positive by *Fasciola* qPCR, but did not have *F. hepatica* reads in tremabiome, despite the qPCR demonstrating no specificity issues towards *C. daubneyi* during assay validation.

**Table 4:**
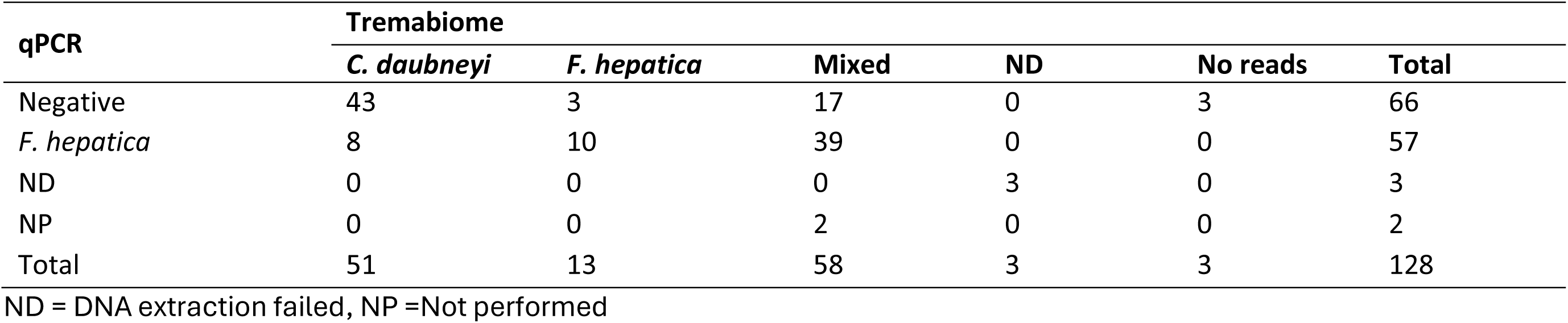
Comparison of qPCR and tremabiome on 128 fluke egg-positive samples.

| qPCR | Tremabiome |  |  |  |  |  |
| --- | --- | --- | --- | --- | --- | --- |
|  | *C. daubneyi* | *F. hepatica* | Mixed | ND | No reads | Total |
| Negative | 43 | 3 | 17 | 0 | 3 | 66 |
| *F. hepatica* | 8 | 10 | 39 | 0 | 0 | 57 |
| ND | 0 | 0 | 0 | 3 | 0 | 3 |
| NP | 0 | 0 | 2 | 0 | 0 | 2 |
| Total | 51 | 13 | 58 | 3 | 3 | 128 |

ND = DNA extraction failed, NP =Not performed

**Table 5:** Comparison of Sanger sequencing and tremabiome on selected samples.

| Tremabiome | Sanger sequencing using ITS2 |  |  |  |  |  | Total |
| --- | --- | --- | --- | --- | --- | --- | --- |
|  | *F. hepatica* | *C. daubneyi* | *Paramphistomum epiclitum* | PSQ | Not performed | ND<sup>1</sup> |  |
| *F. hepatica* | 2 | 0 | 1 | 2 | 8 | 0 | 13 |
| *C. daubneyi* | 0 | 29 | 0 | 9 | 13 | 0 | 51 |
| Mixed | 5 | 14 | 0 | 10 | 29 | 0 | 58 |
| No reads | 0 | 0 | 0 | 3 | 0 | 0 | 3 |
| ND | 0 | 0 | 0 | 0 | 0 | 3 | 3 |
| Total | 7 | 43 | 1 | 24 | 50 | 3 | 128 |

<sup>1</sup>ND = DNA extraction failed; PSQ = poor sequence quality

### Comparison of Sanger sequencing and tremabiome deep amplicon sequencing for species identification

Sanger sequencing identified 7 samples as *F. hepatica* of which tremabiome confirmed two as *F. hepatica*, and five as mixed infections, and none as *C. daubneyi*. Among the 43 samples identified as *C. daubneyi* by Sanger sequencing, tremabiome confirmed 29 as *C. daubneyi* and 14 as mixed infections. Notably, the sample identified as *P. epiclitum* by Sanger sequencing was detected as *F. hepatica* in tremabiome sequencng (Table. 5, Table S5).

## Discussion

In this study we present new approaches for detecting and identifying fluke species in faecal samples in the form of qPCR with high analytical sensitivity and specificity for the detection of *Fasciola* spp. and a deep amplicon sequencing assay which can accurately identify and differentiate between closely related fluke species, such as *F. hepatica*, *F. gigantica*, and *C. daubneyi*. These methods overcome important limitations of microscopic egg examination. We selected the ITS2 and mt-ND1 genetic markers based on their previous application in fluke species identification and their potential to differentiate between closely related flukes [32,39].

Multiple direct and indirect diagnostic approaches are available for diagnosing fluke infections, each with its own set of limitations. However, farmers still require highly sensitive, specific, and cost-effective early diagnostic procedures. Historically, the most widely used traditional direct identification method is detecting fluke eggs in the hosts faeces. This can be accomplished through various techniques such as FLOTAC [46], sedimentation, Flukefinder, or the Kato-Katz thick smear method [47]. Whilst the use of fluke egg counts is simple, however, this diagnostic route can be unreliable due to low intermittent egg deposition in the faeces [48], and it is time-consuming and requires highly trained laboratory staff [37].

Progress in the development of molecular-level diagnostic methods based on DNA detection is underway, including PCR techniques [23], qPCR [22] and the LAMP approach [24]. High throughput deep amplicon sequencing approach of metabarcoded DNA from parasite populations using the Illumina MiSeq platform can offer a low-cost and potentially more accurate alternative to traditional microscopic methods. For instance, adult *Fasciola* spp. and *C. daubneyi* flukes have previously been detected using tremabiome deep amplicon sequencing [32–34] but this method has not been applied to the detection of eggs in faecal samples.

In the present study, species identification through microscopic examination was first checked by PCR followed by Sanger sequencing. Our findings demonstrated that microscopy has limitations in accurately identifying fluke species compared to Sanger sequencing and qPCR and is a time-consuming process, as previously reported by Calvani et al, (2017) [37]. Only about half of the microscopically positive samples (n=51) were confirmed by ITS2 Sanger sequencing to contain *F. hepatica* or C*. daubneyi* DNA. A major contributing factor to this finding could be the presence of mixed infections since fluke eggs are morphologically similar, and this was confirmed by mt-ND1 Sanger sequencing [18,19]. PCR bands were observed on the agarose gel for most of the samples, which indicated successful DNA amplification. However, Sanger sequencing produced poor-quality sequence reads for many samples. Poor sequencing quality can be due to non-specific amplifications, artefacts, and samples containing mixed DNA templates from double infections [49]. Since DNA was extracted from faecal material, additional factors such as low DNA concentration may also have contributed to low-quality Sanger sequencing results. These findings indicated the need for more sensitive and specific molecular diagnostic tools, such as qPCR and tremabiome deep amplicon sequencing, to improve detection accuracy, particularly in complex natural infections.

We repurposed mt-ND1 markers to develop a SYBR Green qPCR assay to detect *Fasciola* spp. The choice to use SYBR Green over fluorescence probe-based systems was due to its cost-effectiveness and simplicity. In contrast, previous studies used TaqMan probes to identify *Fasciola* species [22,37,50]. Our assay’s analytical sensitivity (19.2 fg for *F. hepatica* and 6.4 fg for *F. gigantica* DNA) is lower than that reported in previous studies. For instance, Shi et al, (2020) achieved a detection limit of 1.67 pg of DNA using a SYBR Green qPCR assay targeting the ITS2 region [51]. Similarly, previous studies have demonstrated the ability to detect *F. hepatica* at levels below 10 eggs per gram directly from 150 mg of faecal material using a TaqMan qPCR assay [37] and sensitivities as low as 1 pg/μL [22] and 1.6 pg/μL when targeting the ITS1 region [52]. *F. hepatica* eDNA (14-50 fg) was detected in water samples with similar sensitivity to our assay [53]. In 2024, a qPCR assay was reported which could detect 10 fg of *Fasciola* DNA in water and 1 pg in human stool samples [54]. A limitation of our study is that qPCR was only performed on fluke egg-positive samples due to limited resources. Thus, it was not possible to formally determine diagnostic performance. This will be determined in future work. The diagnostic sensitivity of other qPCR methods was reported to be 66% in human stool samples [52] and 91–100% in sheep and cattle compared to microscopy-based techniques [37]. These estimates assume, however, that microscopy is the gold standard, whereas it could miss positive cases that are detectable using molecular methods.

A limitation of our qPCR assay is that we are only able to detect *Fasciola* in DNA extracted from sedimented material, not DNA extracted directly from faecal samples. Raw faecal samples were tested in this study using the same DNA extraction methodology as sedimented eggs, but it was not possible to detect *Fasciola* DNA using the qPCR methodology described in the study (data not shown). However, sensitivity improved when *Fasciola* eggs were first concentrated using faecal egg sedimentation before applying the bead-beating approach for DNA isolation [37]. Similarly, other studies also employed molecular procedures following the sedimentation process [18,50,55], and a few studies have applied molecular techniques to detect natural *Fasciola* infections directly from faecal material and reported limitations [37,55]. One possible approach is LAMP [56], as this has demonstrated low detection limits for *Fasciola* spp. DNA and results can be observed with the naked eye [24,25]. A recent study successfully detected *F. hepatica* in DNA extracted directly from faeces with a commercial kit, using both LAMP and PCR methodology targeting ITS2 region [57]. Further work is needed to simplify extraction protocols.

In ruminants, fluke species often occur in complex and overlapping infections; for instance, *F. hepatica* and *C. daubneyi* co-infections have been observed in cattle and sheep in the UK [58,59] and the same has been reported elsewhere in Europe [18,60], including in Ireland [61] and Germany [62]. We found that the microscopic egg identification for closely related flukes, such as *F. hepatica* and *C. daubneyi*, is challenging due to their similar size and shape. Therefore, the tremabiome technology was developed utilising universal ITS2 rDNA markers to differentiate between multiple fluke species. The techniques were validated using fluke egg DNA, isolated from faecal samples. To our knowledge, this is first time the tremabiome deep amplicon sequencing approach has been applied to detect mixed fluke infections in faeces. Previously the approach was applied to detect adult fluke samples [33,34].

The tremabiome technique generated sequence reads for *F. hepatica*, *F. gigantica*, and *C. daubneyi*. However, the proportion of sequence reads deviated from expected percentages. For instance, we evaluated the assay’s ability to accurately determine the relative species proportions in pairwise combinations of *F. hepatica* and *C. daubneyi*. The results consistently showed a higher proportion of reads for *C. daubneyi* compared to *F. hepatica* across all mixtures. Such variation may arise from factors including the primers used for the target loci, conserved priming sites, variations in DNA template concentrations during sample handling, the number of PCR cycles, and the copy number of the target DNA locus [63]. Previously, for nematodes, species-specific representation biases were addressed by calculating correction factors using L3 larval population DNA from different nematode species [29]. In the present study, while working with fluke egg DNA obtained via faecal sedimentation, calculation of correction factors did not remove sequence biases (File S5). This might be due to differences in the eggshell chemistry or stability hardness of eggshells that has been described between *F. hepatica* and *C. daubneyi* [64], leading to variations in DNA extraction efficiency. However, we employed mechanical disruption before DNA isolation to mitigate this issue [37]. Additionally, we used universal ITS2 primers to detect multiple fluke species in a single deep amplicon sequencing run. Using species-specific primers in deep amplicon sequencing could be a potential solution to reduce sequence biases [65]. Further, bioinformatics analysis of sequencing data might introduce biases, leading to inaccuracies in species proportion estimations. One major challenge can be the limited availability of reference sequence reads in the NCBI database, for example, for *F. hepatica* 50 unique sequences and for *C. daubneyi* 19 unique sequences were found, which can impact species specific reads identification. Additionally, taxa represented by low numbers of sequencing reads may pose a problem, as these low-frequency reads were removed during data filtering while eliminating artefacts, resulting in the underrepresentation of actual sequence reads. Therefore, there is a need for more reference sequence data, which can enhance our capability of accurately distinguishing true sequences. When the tremabiome technology was applied to field samples, many *F. hepatica* and *C. daubneyi* co-infections were identified. Since *F. hepatica* is more pathogenic and economically detrimental [6,66] than *C. daubneyi* [67,68], and treatment choices differ, our method provides a valuable tool for differentiating co-infections of these two significant parasites using faecal egg samples.

When applying the tremabiome approach to field samples, just over half of the microscopically positive *F. hepatica* samples were confirmed by tremabiome deep amplicon sequencing. Moreover, there was a significant correlation between the identification of *F. hepatica* infections using qPCR and the tremabiome approach. Notably, the tremabiome approach generated *F. hepatica* sequence reads in samples which tested negative by qPCR. Conversely, tremabiome deep amplicon sequencing did not produce *F. hepatica* reads for a few qPCR-positive samples. It has been reported that false negative results for molecular tests may be due to low egg count in faecal samples [18]. Furthermore, one sample identified as *P. epiclitum* by Sanger sequencing was not confirmed by tremabiome, which instead identified it as *F. hepatica*. Previously *P. leydeni* has been reported in sheep [69], and deer [70] in Ireland, but was not identified in the UK in our study. Therefore, each method has limitations, but tremabiome remained the preferred tool for species-level fluke identification as it can identify mixed infections.

Regarding the implementation of these methods, microscopy can be a suitable option for analysing a small number of samples due to its low cost and accessibility, although, there were potential issues of misidentifications [18,37]. In particular, a high proportion of samples identified as positive for *F. hepatica* based on egg morphology were confirmed by tremabiome to contain *C. daubneyi*, despite careful identification. For medium to high sample volumes, qPCR is suitable to identify *Fasciola* spp. infections only and tremabiome deep amplicon sequencing is potentially more effective choices for species level differentiation of different flukes with an advantage of high throughput, as a single Illumina MiSeq run can process up to 384 samples simultaneously. This capability makes the method suitable for both research and diagnostic applications. Presently, this study offers evidence of the high prevalence of *F. hepatica* and *C. daubneyi* in UK ruminants. Expanding this method to larger sample sizes across the UK and in other countries would provide a more comprehensive epidemiological understanding of these infections.

The sequencing data generated from the set of natural field samples enhanced our understanding of the genetic variation within fluke populations by revealing their ASVs. Notably with ITS2 markers, we observed more ASVs for *F. hepatica* than *C. daubneyi*, indicating possible greater genetic diversity within *F. hepatica* populations. Previously, *Fasciola* species in Pakistan were differentiated using ITS2 markers in adult worms [32,39], while high genetic diversity was reported using mt-ND1 markers [32,33]. Similarly, a study from Spain and Peru identified *Fasciola* flukes using nuclear DNA markers but reported high genetic diversity based on mitochondrial markers [71]. Further, high genetic diversity and gene flow among *Fasciola* populations have been reported in the UK, using microsatellite markers [72,73]. Therefore, the ITS2 rDNA provides a useful taxonomic resolution [32,39], however, mitochondrial markers such as (ND1 and COX1) and genetic markers are typically preferred for detailed population genetic studies [33,71,72,74–76]. Therefore, further investigations using mitochondrial ND1 and nuclear genetic markers are required to understand the genetic diversity of these fluke populations and is work in progress.

Although validated on cattle and sheep samples in this study, both the qPCR and tremabiome methods have a strong potential for the *Fasciola* spp. detection in humans, particularly in endemic regions where there is a possibility of zoonotic transmission of *F. hepatica* and *F. gigantica*. Application of these methods on human faecal samples could improve case detection, fluke species identification, and epidemiological understanding.

## Conclusion

In conclusion, this study presents the first use of tremabiome deep amplicon sequencing for detecting mixed infections of *F. hepatica* and *C. daubneyi*, and provides a direct comparison between microscopy, PCR, Sanger sequencing, qPCR and tremabiome methods for differentiating between fluke species. The tremabiome approach was highly effective for detecting mixed fluke infections, compared to other techniques utilised in this study, demonstrating high frequency of *C. daubneyi* and *F. hepatica* co-infections in farmed ruminants in the UK. The methods were primarily validated using samples from natural infections, with DNA extracted from faecal sedimented eggs, which allowed an easy and non-invasive sampling approach at the farm level. Tremabiome and qPCR are promising tools to complement microscopy in fluke disease surveillance and control in livestock and humans.

## Supporting information

<u>File S1</u> **Relative proportions of *F. hepatica* and *C. daubneyi* in egg DNA mixtures** (XLSX)

<u>File S2</u> **Mock egg mixtures with gradually decreasing counts of *F. hepatica* eggs** (XLSX)

<u>File S3</u> **Mock egg mixtures with gradually decreasing counts of *C. daubneyi* eggs** (XLSX)

<u>File S4</u> **Deep amplicon sequence reads generated from field samples from cattle and sheep** (XLSX)

<u>File S5</u> **Correction factors calculations** (PDF)

<u>Table S1</u> **Sample information** (XLSX)

<u>Table S2</u> **Reference sequences downloaded from NCBI and unique sequence count** (XLSX)

<u>Table S3</u> **mt-ND1 and ITS2 primer sequences** (PDF)

<u>Table S4</u> **Genetic distances for different fluke species based on ITS2 marker** (XLSX)

<u>Table S5</u> **Sample (n=128) details used for comparison of techniques for microscopy, PCR, Sanger sequencing, qPCR, and tremabiome** (XLSX)

<u>Table S6</u> **Coefficients of variation for qPCR** (PDF)

<u>Fig. S1 and Fig. S2</u> **Analytical sensitivity and specificity of qPCR** (PDF)

## Supporting information

File S1 Relative proportions of F. hepatica and C. daubneyi in egg DNA mixtures (XLSX)

File S2 Mock egg mixtures with gradually decreasing counts of F. hepatica eggs (XLSX)

File S3 Mock egg mixtures with gradually decreasing counts of C. daubneyi eggs (XLSX)

File S4 Deep amplicon sequence reads generated from field samples from cattle and sheep (XLSX)

File S5 Correction factors calculations (PDF)

Table S1 Sample information (XLSX)

Table S2 Reference sequences downloaded from NCBI and unique sequence count (XLSX)

Table S3 mt-ND1 and ITS2 primer sequences (PDF)

Table S4 Genetic distances for different fluke species based on ITS2 marker (XLSX)

Table S5 Sample (n=128) details used for comparison of techniques for microscopy, PCR, Sanger sequencing, qPCR, and tremabiome (XLSX)

Table S6 Coefficients of variation for qPCR (PDF)

Fig. S1 and Fig. S2 Analytical sensitivity and specificity of qPCR (PDF)

## Acknowledgements

Part of this work was carried out using the computational HPC facilities and support provided by the Research Computing Services team within IT Services at University of Surrey, specifically the Eureka2 HPC cluster (https://docs.pages.surrey.ac.uk/research_computing/hpc/clusters/eureka2.html).

This research was funded in whole, or in part, by the UK Research and Innovation (UKRI), Biotechnology and Biological Sciences Research Council (BBSRC) through the FoodBioSystems Doctoral Training Programme (BB/T008776/1) and the Sir Halley Stewart Trust (3153). For the purpose of Open Access, the authors have applied a Creative Commons Attribution (CC BY) public copyright licence to any Author Accepted Manuscript version arising from this submission.

## Credit authorship contribution statement

Muhammad Abbas: conceptualisation, investigation, methodology, bioinformatics, validation, visualisation, data curations and analysis, writing original draft, review and editing; Kezia Kozel: investigation, methodology, formal analysis, validation, writing original draft, review and editing; Olukayode Daramola: writing original draft, review, formal analysis, data curation, supervision; Nick Selemetas: review and editing, supervision; Qasim Ali: resources; Shoaib Ashraf: resources; Isah Ibrahim: resources; Inaki Deza-Cruz: resources; review and editing; Sai Fingerhood: resources; review and editing; Mark W Robinson: resources; review and editing; Eric R Morgan: funding acquisition; supervision, review and editing; Umer Chaudhry: conceptualisation, formal analysis, validation, data curation, writing, review and editing, supervision; Martha Betson: conceptualisation, formal analysis, writing review and editing, supervision, funding acquisition, project administration.

## Data availability

All sequencing data produced in the paper are available at NCBI BioProject ID PRJNA1273189, and accession numbers PV752160-PV752182, PV752186-PV752205, PV752431-PV752462, PV752238-PV752240, PV752248-PV752250, PV752375-PV752429, PV752270.

In addition, sequence data, scripts and codes are available at Mendeley data base DOI: 10.17632/zyvwc6ppy8.1.

All other data are reported in the paper and associated supplementary material.

