## Supplementary material for "Development of a qPCR assay and tremabiome deep amplicon sequencing method for differentiation of fluke species in livestock": File S5 Correction factors calculations (PDF)

Accurately quantifying species abundance using deep amplicon sequencing approaches remains challenging due to sequence biases introduced during PCR amplification [1]. The number of sequence reads generated for different species using a single universal primer pair may vary due to the possibility of mismatches in primer binding regions [2,3]. Additionally, factors such as the target sequence itself can introduce amplification bias. For example, in ribosomal DNA amplicon mixtures of variable lengths, shorter sequences are preferentially amplified, and GC content can also influence amplification efficiency. Moreover, variations in target locus copy number among species can affect abundance estimates from PCR-based methods [4]. In addition, the efficiency of the DNA extraction method and species-specific variations in the amplification of target fragments could be the contributing factors [5,6].

To address this, we examined sequence data from various prepared mock mixes of DNA from different fluke eggs that were subjected to PCR cycles 25x, 30x, and 35x runs. As illustrated in Fig. 1, biases were observed in species representation across different cycle numbers in mock mixes made from fluke egg DNA. These biases may be due to multiple factors described above.

We calculated correction factors for each species by dividing the true proportion of species in the pool (determined from egg count percentages) by the average proportion of deep amplicon sequences obtained from the assay in triplicates (correction factor = %Actual / %Observed). This correction factor helped reduce sequencing biases, aligning the data more closely with expected values (Fig. 1). Correction factors were determined for three fluke species using egg DNA, based on the available laboratory resources. On the one hand, Fig. 1 applies these correction factors to bring species proportions closer to their expected values. On the other hand, Fig. 2 and 3 (Files S1, S2 and S3) indicated that the correction factors reduced sequencing biases, but most of the biases remained statistically significant. The corresponding figure legends provide specific details of the statistical tests performed.

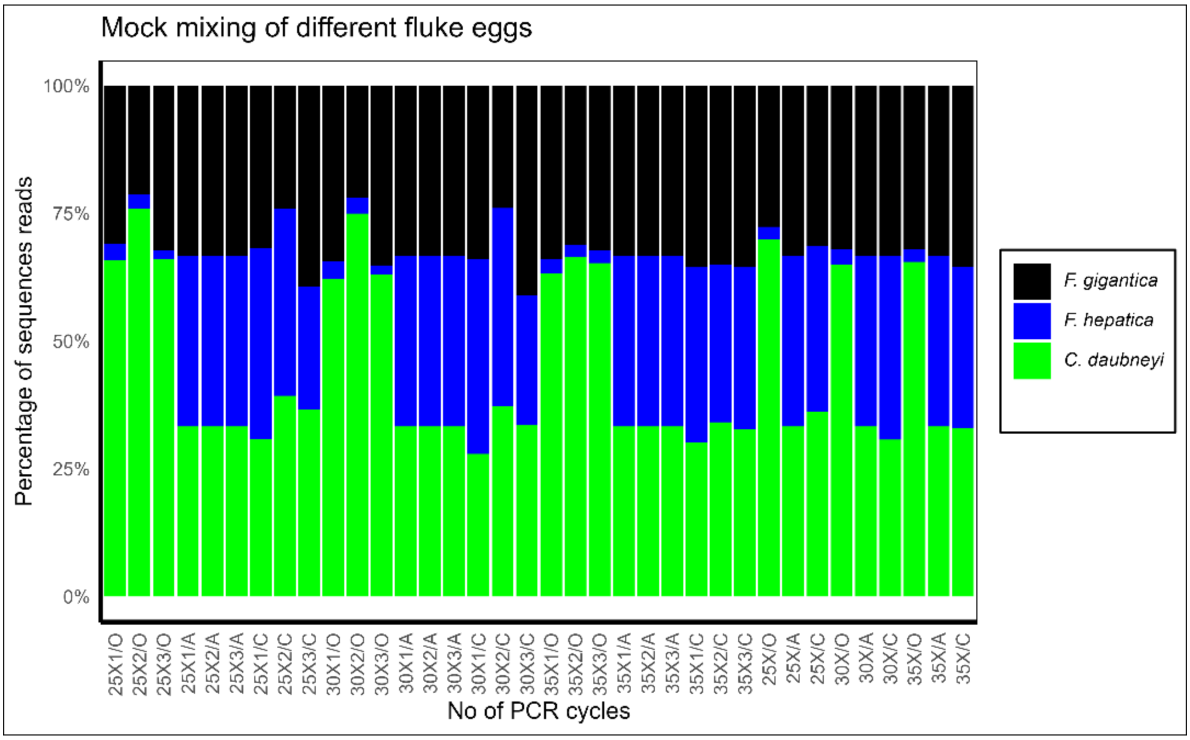


**Fig. 1:** Correction factors and sequence representation bias in deep amplicon sequencing of a mock mixture of three fluke species: *F. hepatica*, *F. gigantica*, and *C. daubneyi*. DNA was extracted in triplicate from pooled samples containing 250 eggs of each species. The DNA mixture was amplified using PCR at three cycle levels (25X, 30X, and 35X), with triplicate testing for each pool. The x-axis indicates PCR cycle numbers, while the y-axis represents the percentage of ITS-2 rDNA sequence reads for each species, categorised as observed (O), actual (A), and corrected (C). A non-parametric chi-square test revealed significant differences between observed ITS2 sequence reads and actual proportions of egg pools at all cycle levels (25X, 30X, and 35X). The *p*-values calculated at 25X PCR cycles, ($p$=1.29×10$^{-13}$, $p$=1.37× 10$^{-19}$, $p$=3.25×10$^{-14}$), at 30X PCR cycles ($p$=5.12×10$^{-12}$, $p$=8.24×10$^{-19}$ and $p$=6.5×10$^{-13}$) and 35X PCR cycles ($p$=1.35×10$^{-12}$, $p$=3,67×10$^{-14}$ and $p$=1.35×10$^{-13}$). Correction factors were applied (*F. gigantica* = 1.09, *F. hepatica* = 12.67, and *C. daubneyi* = 0.49), significantly reducing bias in the data. The non-parametric Chi-square test on corrected reads showed 25X ($p$=0.697, $p$=0.14, and $p$=0.138), 30X ($p$=0.454, $p$=0.135, and $p$=0.156), and 35X ($p$=0.785, $p$=0.853, and $p$=0.886). Further, one-way ANOVA confirmed that the number of PCR cycles (25X, 30X, and 35X) did not affect the proportional representation of *F. gigantica* ($p$ = 0.276), *F. hepatica* ($p$ = 0.993), and *C. daubneyi* ($p$ = 0.975). These findings show the effectiveness of deep amplicon sequencing in identifying species representation across PCR cycle levels.

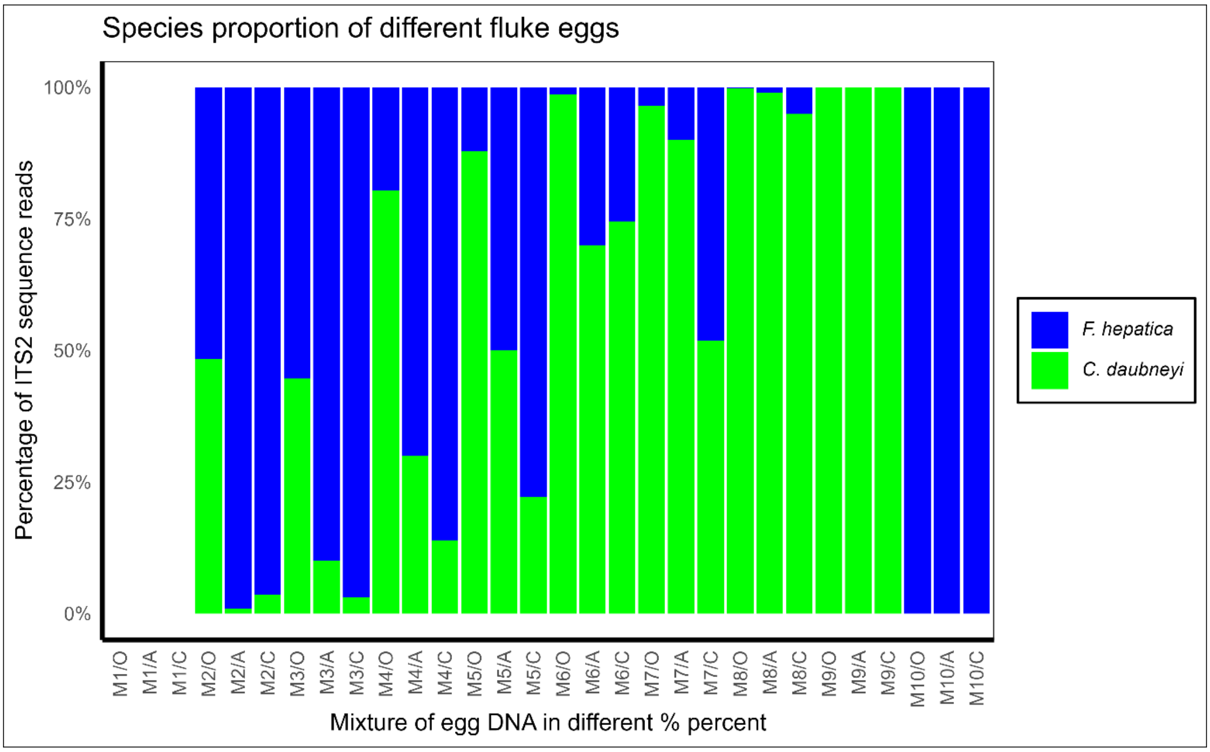


**Fig. 2:** Relative proportions of *F. hepatica* and *C. daubneyi* based on egg DNA mixtures, assessed using a deep amplicon sequencing assay. DNA was extracted in triplicate from mock pools containing varying ratios of these two fluke species, enabling evaluation of the assay's accuracy across a range of species proportions. The x-axis represents egg mixtures with *F. hepatica*: *C. daubneyi* ratios organised as: M1 (negative control), M2 (99:1), M3 (90:10), M4 (70:30), M5 (50:50), M6 (30:70), M7 (10:90), M8 (1:99), M9 (100% *C. daubneyi*), and M10 (100% *F. hepatica*). Results are further categorised into observed (O), actual (A), and corrected (C) values. The y-axis shows the percentage of ITS-2 rDNA sequence reads for each species. Chi-square test results for uncorrected reads vs. expected proportions indicate significant deviations ($p < 0.05$) for most mixtures, with the highest significance of biases observed in M2 (99:1) and M3 (90:10) ($p$=0, $p$=6.79 × $10^{-31}$, respectively). Minor biases were noted in M7 (10:90) ($p$ =0.031), while no significant difference was found in M8 (1:99) ($p$=0.428). These results highlight the varying accuracy of uncorrected reads depending on species proportions. Further, the chi-square test was applied to corrected reads vs. expected proportions across egg mixtures. Significant deviations were observed for M2 ($p$=0.0102), M3 ($p$=0.0212), M4 ($p$=0.0004), M5 ($p$=2.56 × $10^{-8}$), M7 ($p$=5.39 × $10^{-37}$), and M8 ($p$=3.75 × $10^{-5}$). No significant difference was found for M6 ($p$=0.3262). Therefore, the correction factor did not resolve the observed sequence biases in most mixtures.

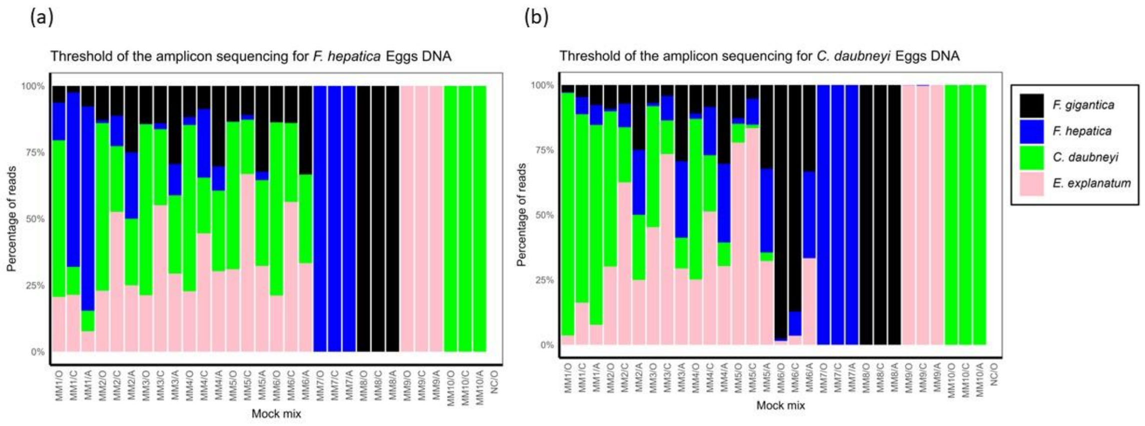


**Fig. 3**: Illustrates the detection threshold of the amplicon sequencing assay with progressively lower counts of *F. hepatica* and *C. daubneyi* eggs. Mock pools were created for each of the available four fluke species (*F. hepatica*, *F. gigantica*, *C. daubneyi*, *E. explanatum*), with separate pools for (a) *F. hepatica* (b) *C. daubneyi* with decreasing egg numbers: 500, 50, 20, 15, 5, and 0 eggs. The mock mixes are from MM1 to MM6, while MM7 to MM10 represent sequencing reads from individual species derived from egg DNA. The x-axis indicates DNA mixes used in the assay categorised as observed (O), actual (A), and corrected (C), while the y-axis represents the percentage of ITS-2 rDNA sequence reads for each species. These results demonstrate the assay's sensitivity, with detectable reads down to 5 eggs for both *F. hepatica* and *C. daubneyi*. (a) Performance of a chi-square test on *F. hepatica* uncorrected reads revealed significant deviations between observed and actual values across all mixes, with *p*-values MM1 $p$=1.85 × $10^{-89}$, MM2 $p$=1.04 × $10^{-18}$, MM3 $p$=1.30 × $10^{-13}$, MM4 $p$=3.51 × $10^{-11}$, MM5 $p$=9.92 × $10^{-7}$, and MM6 $p$=4.68 × $10^{-10}$. For corrected reads, the test showed reduced deviations, reflecting the impact of corrections in minimising discrepancies, however, significant differences between corrected and actual values noted across the mixes with *P* values for MM1 $p$=9.42 × $10^{-7}$, MM2 $p$= 7.41 × $10^{-10}$, MM3 $p$ =2.47 × $10^{-8}$, MM4 $p$=5.35 × $10^{-12}$, MM5 $p$=3.85 × $10^{-12}$, and MM6 $p$=4.38 × $10^{-6}$. (b) The chi-square test for *C. daubneyi* uncorrected reads revealed significant differences between observed and actual values across all mixes. The *p*-values were MM1 $p$=0.0011, MM2 $p$=9.20×$10^{-18}$, MM3 $p$=1.67×$10^{-33}$, MM4 $p$=1.15×$10^{-74}$, MM5 $p$=1.60×$10^{-23}$ and MM6 $p$=4.57×$10^{-40}$. Further, there were significant differences between corrected and actual values across all mixes. The *p*-values calculated were for MM1 *P*=0.0116, MM2 $p$=3.52×$10^{-17}$, MM3 $p$=8.61×$10^{-22}$, MM4 $p$=2.90×$10^{-11}$, MM5 $p$=6.85×$10^{-26}$ and MM6 $p$=2.97×$10^{-28}$. These results indicate that the corrections reduced the level of deviation compared to uncorrected reads, but the differences remained statistically significant across all mixes.
