## Supplementary material for "Development of a qPCR assay and tremabiome deep amplicon sequencing method for differentiation of fluke species in livestock": Table S3 mt-ND1 and ITS2 primer sequences (PDF)

mt-ND1 and ITS2 primer sequences for amplifying mitochondrial region NADH dehydrogenase 1 and internal transcribed spacer 2 region of the rDNA respectively. The adaptor sequences are in italics, Forward and reverse primers are underlined. The 'N's represent random nucleotides added between the Illumina adaptor sequences and the locus-specific primers. Modified phosphate bonds at positions indicated with asterisks

| Primer ID | Target | Primer sequence | Product length (bps) | Remarks | Reference |
|---|---|---|---|---|---|
| mt-ND1 | Mitochondrial markers | UN2 (For): 5'-GTTTAAGTTTGTGTTTTTTC-3'<br>ND1 (Rev): 5'-CACCATAACTCCCCCAAACCA-3' | 311 | Primers used in qPCR to amplify mitochondrial region NADH dehydrogenase 1 | (Rehman et al., 2020) |
| ITS2 | Coding regions of 5.8S and 28S rDNA | ITS2 (For): 5'-GGTGGATCACTCGGCTCGTG-3'<br>ITS2 (Rev): 5'-TTCCTCCGCTTAGTGATATGC-3' | 490-743 | Universal ITS2 region primers | (Chaudhry et al. 2016) |
| AD_For ITS2 | ITS2 + adaptor primer | *TCGTCGGCAGCGTCAGATGTGTATAAGAGACAG*<br>GGTGGATCACTCGGCTCG*T*G | 490-743 | Forward direction | This work |
| AD_For 1N ITS2 | ITS2 + adaptor primer | *TCGTCGGCAGCGTCAGATGTGTATAAGAGACAGN*<br>GGTGGATCACTCGGCTCG*T*G | 490-743 | Forward direction | This work |
| AD_For 2N ITS2 | ITS2 + adaptor primer | *TCGTCGGCAGCGTCAGATGTGTATAAGAGACAGNN*<br>GGTGGATCACTCGGCTCG*T*G | 490-743 | Forward direction | This work |
| AD_For 3N ITS2 | ITS2 + adaptor primer | *TCGTCGGCAGCGTCAGATGTGTATAAGAGACAGNNN*<br>GGTGGATCACTCGGCTCG*T*G | 490-743 | Forward direction | This work |
| AD_Rev ITS2 | ITS2 + adaptor primer | *GTCTCGTGGGCTCGGAGATGTGTATAAGAGACAG*<br>TTCCTCCGCTTAGTGATAT*G*C | 490-743 | Reverse direction | This work |
| AD_Rev 1N ITS2 | ITS2 + adaptor primer | *GTCTCGTGGGCTCGGAGATGTGTATAAGAGACAGN*<br>TTCCTCCGCTTAGTGATAT*G*C | 490-743 | Reverse direction | This work |
| AD_Rev 2N ITS2 | ITS2 + adaptor primer | *GTCTCGTGGGCTCGGAGATGTGTATAAGAGACAGNN*<br>TTCCTCCGCTTAGTGATAT*G*C | 490-743 | Reverse direction | This work |
| AD_Rev 3N ITS2 | ITS2 + adaptor primer | *GTCTCGTGGGCTCGGAGATGTGTATAAGAGACAGNNN*<br>TTCCTCCGCTTAGTGATAT*G*C | 490-743 | Reverse direction | This work |
