## Supplementary material for "Development of a qPCR assay and tremabiome deep amplicon sequencing method for differentiation of fluke species in livestock": Table S6 Coefficients of variation for qPCR (PDF)

**Table S6:** Variation in *F. hepatica and F. gigantica* SYBR green qPCR Cq values within and between assays using data from analytical sensitivity test

| | Cq values observed for *F hepatica* DNA | | | | Cq values observed for *F. gigantica* DNA | | |
|---|---|---|---|---|---|---|---|
| DNA conc. | Mean | SD | CV% | DNA conc. | Mean | SD | CV% |
| 300 pg | 21.96 | 0.16 | 0.72 | 500 pg | 18.62 | 0.147 | 0.78 |
| 60 pg | 24.73 | 0.327 | 1.32 | 100 pg | 21.13 | 0.066 | 0.31 |
| 12 pg | 28.59 | 1.717 | 6.00 | 20 pg | 24.16 | 0.195 | 0.80 |
| 2.4 pg | 30.84 | 0.347 | 1.12 | 4 pg | 26.85 | 0.023 | 0.08 |
| 480 fg | 34.46 | 0.330 | 0.95 | 800 fg | 29.94 | 0.142 | 0.47 |
| 96 fg | 34.43 | 0.602 | 1.74 | 160 fg | 32.91 | 0.29 | 0.88 |
| 19.2 fg | 38.26 | 0.343 | 0.89 | 32fg | 35.73 | 0.614 | 1.71 |
| | | | | 6.4 fg | 38.24 | 1.063 | 2.77 |

Cq: PCR cycle number, SD: standard deviation, CV: coefficient of variation
