## Supplementary material for "Development of a qPCR assay and tremabiome deep amplicon sequencing method for differentiation of fluke species in livestock": Fig. S1 and Fig. S2 Analytical sensitivity and specificity of qPCR (PDF)

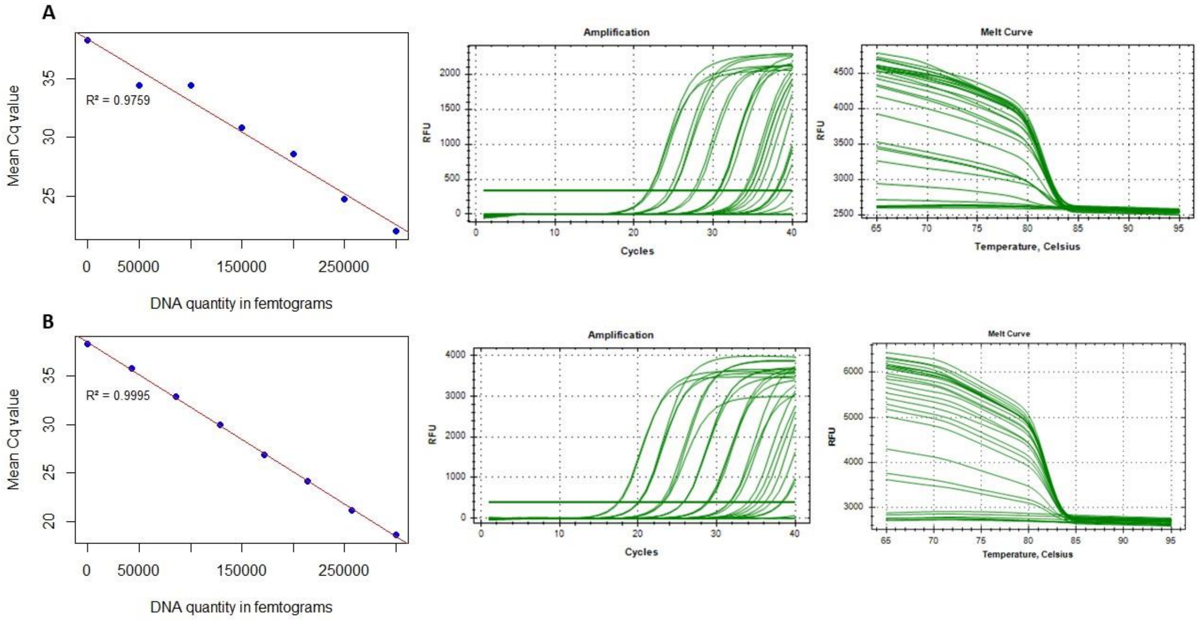


**Fig. S1:** Sensitivity of qPCR targeting mt-ND1 mitochondrial DNA of (A) *F. hepatica* and (B) *F. gigantica*. The standard curve was plotted using qPCR data from 5-fold serial dilutions of adult worm DNA extracted from the head of *F. hepatica* and *F. gigantica*. DNA quantification was performed via Qubit and verified in the qPCR assay. The average of three Cq values was plotted against the DNA quantity to create the standard curve. The melt curves for *F. hepatica* and *F. gigantica* at 81.5°C are also shown.

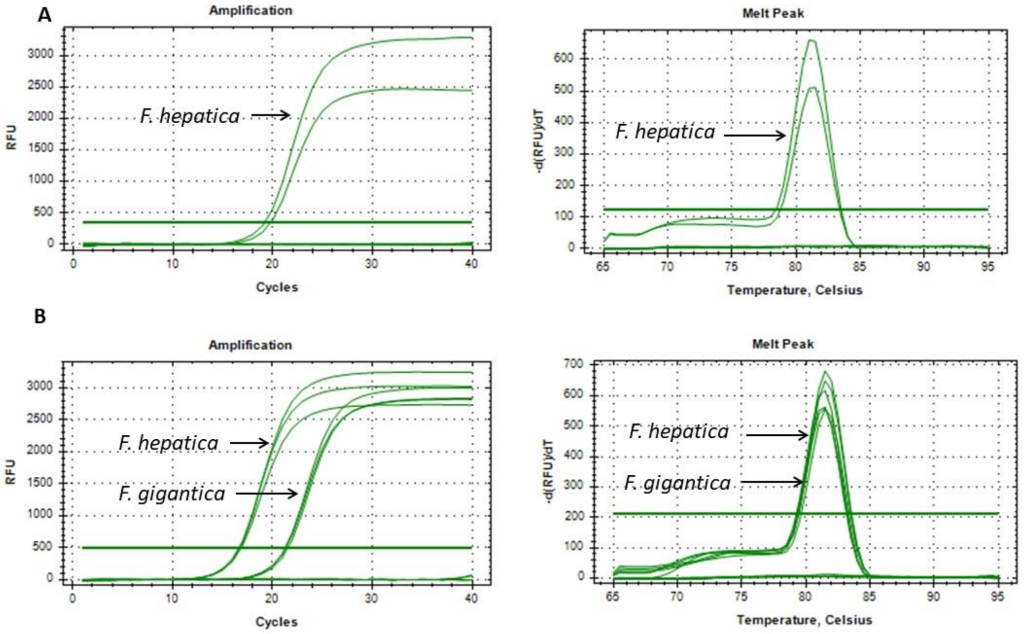


**Fig. S2:** Specificity of mt-ND1 primers in qPCR showing amplification of *F. hepatica* and *F. gigantica*. (A) Specificity of mt-ND1 primers evaluated with DNA of other flukes *Paramphistomum*, *E.* e*xplanatum*, *C. daubneyi.* (B) Specificity of mitochondrial primers tested with DNA of nematodes, *T. circumcincta*, *Ascaris*, and *Trichuris*. Amplification curves are shown on the left-hand side and melt curves on the right-hand side. There is no cross-amplification with non-target species in any of the reactions.
